# The oxytocin-modulated brain circuit that synchronizes heart rate with breathing

**DOI:** 10.1101/2023.09.26.559512

**Authors:** Julie Buron, Ambre Linossier, Christian Gestreau, Fabienne Schaller, Roman Tyzio, Marie-Solenne Felix, Valéry Matarazzo, Muriel Thoby-Brisson, Françoise Muscatelli, Clément Menuet

**Affiliations:** INMED, INSERM, Aix-Marseille University, Marseille, France; INCIA, CNRS, Université de Bordeaux, Bordeaux, France

## Abstract

The variation in heart rate in phase with breathing, called respiratory sinus arrhythmia (RSA), is cardio-protective^1,2^. RSA amplitude provides an index of health and physical fitness used both clinically, and by the broader population using “smart” watches. Relaxation and positive socio-emotional states can amplify RSA^3^, yet the underlying mechanism remains largely unknown. Here, we identify a hypothalamus-brainstem neuronal network through which the neuromodulator oxytocin (OT), known for its relaxing and prosocial effects^4^, amplifies RSA during calming behavior. OT neurons from the caudal paraventricular nucleus in the hypothalamus were found to regulate the activity of a subgroup of inhibitory neurons in the pre-Bötzinger complex, the brainstem neuronal group that generates the inspiratory rhythm. Specifically, OT amplifies the inspiratory glycinergic input from pre-Bötzinger complex neurons to cardiac-innervating parasympathetic neurons in the nucleus ambiguus. This leads to amplified respiratory modulation of parasympathetic activity to the heart, thereby amplifying RSA. Behaviorally, OT neurons participate in the restoration of RSA amplitude during recovery from stress. This work shows how a central action of OT induces a physiologically beneficial regulation of cardiac activity during a calming behavior, providing a foundation for therapeutic strategies for anxiety disorders and coping with stress. Furthermore, it identifies a phenotypic signature of a subpopulation of neurons controlling RSA, namely pre-Bötzinger complex neurons expressing the OT-receptor, enabling the specific modulation of RSA amplitude to resolve its physiological and psychological functions.

## Main text

Heart rate (HR) is highly variable. It is continuously regulated by the autonomic nervous system to maintain blood gas homeostasis, and to produce physiological manifestations of emotional states such as increased HR during stress and decreased HR during relaxation. A predominant source of HR variability originates from its coupling with respiratory activity, called respiratory sinus arrhythmia (RSA), where HR increases during inspiration and decreases during expiration. Physiologically, RSA optimizes cardiac efficiency, with increased cardiac activity occurring during inspiration when the myocardium receives oxygenated blood and *vice versa* during expiration, while maintaining physiological levels of arterial CO_2_^1^. Yet RSA has broader impact. The amplitude of RSA - i.e. the difference between maximal HR during inspiration and minimal HR during expiration - is larger in young individuals and in well-trained athletes, decreases with age, and is dramatically reduced in hypertension and chronic heart failure^1^. Restoration of RSA amplitude in animal models of chronic heart failure, using a biofeedback heart pacemaker, induces large and persistent improvements in cardiac function^2,5^. RSA amplitude is also strongly modulated by emotional states. RSA is amplified during relaxation and positive socio-emotional states like parental bonding, while it is strongly decreased or even becomes negative in anxious or depressive people, and in autism spectrum disorder^3,6,7^. Interestingly, the neuropeptide oxytocin (OT), widely studied for its relaxing, anxiolytic and pro-social effects^4^, has been found to amplify RSA after intranasal administration in humans^8^.

RSA is mainly generated *via* parasympathetic activity to the heart, mediated by the vagus nerve, which originates from two brainstem nuclei: the nucleus ambiguus (nA) and the dorsal motor nucleus of the vagus nerve (DMV)^9^. Cardiac-innervating nA (nA^Cardiac^) neurons, whose activity decreases HR, are inhibited during inspiration and activated during expiration, thereby generating RSA^9–11^. DMV^Cardiac^ neurons are tonically active, and can regulate mean HR (mHR)^9^. The inhibition of nA^Cardiac^ neuronal activity during inspiration most likely arises from the adjacent pre-Bötzinger Complex (preBötC), the neuronal group that generates the inspiratory rhythm^12–14^. Also, pharmacological studies and the presence of a high density of OT fibers in the preBötC/nA area at the macroscopic level suggest that endogenous OT, which is primarily synthesized by neurons in the paraventricular (PVN) and supraoptic nuclei of the hypothalamus, could modulate the activity of preBötC and/or nA neurons^15–17^. Here, by employing a variety of experimental approaches, we tested whether OT could amplify RSA *via* a central action on preBötC/nA neurons, with the dual aim of specifying the neuronal circuitry that generates RSA, and identifying a mechanism for central RSA amplification with its behavioral relevance.

### PVN^OT^→preBötC/nA neurons induce an amplification of RSA and a decrease in mHR

To determine if preBötC and/or nA neurons receive OT projections, we immunolabelled OT fibers and labelled preBötC or nA neurons in adult WT mice (Fig. 1a, Extended Data Fig. 1a, b). The preBötC is defined by its anatomical location in the ventrolateral medulla oblongata, and its function generating the inspiratory rhythm^18–20^. It contains a network of heterogeneous neurons, with subgroups that are distinguishable by their developmental origin (e.g. Dbx1^+^), neurochemistry (e.g. glutamatergic, glycinergic, GABAergic), receptor expression (e.g. µ-opioid receptor (µO-R), neurokinin 1 receptor (NK1-R)), or projection profile (e.g. projection to the contralateral preBötC to synchronize the generation of the inspiratory rhythm^21^). To locate the preBötC, we used the addition of three identifying criteria: the generation of the inspiratory rhythm, the axonal projections to the contralateral preBötC, and the classical anatomical landmarks for the preBötC (in particular the shape of the inferior olives^22^ and the rostro-caudal position relative to the 4^th^ ventricle). We made extracellular recordings to precisely localize the characteristic pre-inspiratory/inspiratory neurons of the preBötC^12^ (all preBötC injections in this study were guided this way, Extended Data Fig. 1b), and we injected the retrograde neuronal tracer, FluoroGold, unilaterally at this coordinate. The contralateral preBötC was then defined by boundaries surrounding FluoroGold^+^ neurons at the classical preBötC anatomical landmarks (Extended Data Fig. 1b, c). On the other hand, nA neurons, including nA^Cardiac^ neurons, form a nucleus of parasympathetic preganglionic neurons, predominantly sending their axons into the vagus nerve. nA neurons were labeled by intraperitoneal FluoroGold injection (Extended Data Fig. 1a, c). We found dense OT fibers throughout the rostro-caudal extent of the preBötC, some in close apposition to contralaterally-projecting preBötC neurons, but fewer OT fibers near nA neurons (Fig. 1a, Extended Data Fig. 1c, d). To identify which OT neurons project to preBötC/nA neurons, we injected the retrograde neuronal tracer, cholera toxin B, bilaterally into the preBötC/nA (Fig. 1b, Extended Data Fig. 1e-g). OT→preBötC/nA neurons were restricted to the caudal dorso-lateral PVN, forming a dense and specific cluster of PVN^OT^ neurons that project onto preBötC/nA neurons.

**Fig. 1:**
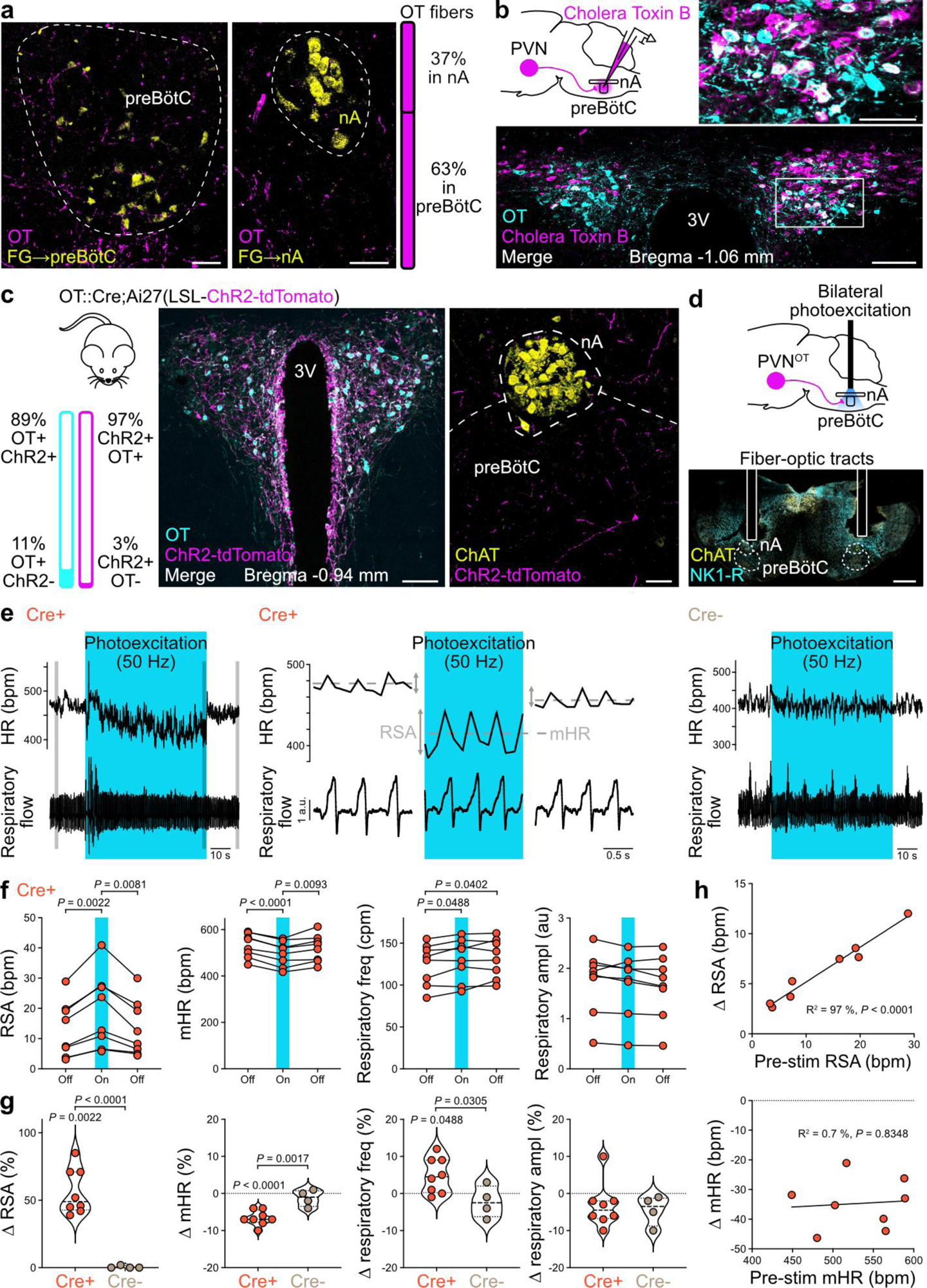
OT fibers from the caudal PVN to the preBötC/nA can amplify RSA, decrease mHR and increase respiratory frequency in adult freely moving mice at rest. **a**, Immunohistochemical labeling of OT fibers in the preBötC and nA. The nA was identified using FluoroGold (FG) retrograde tracing following intraperitoneal injection (Extended Data Fig. 1a). The preBötC was identified using a combination of FG retrograde tracing from injections in the contralateral preBötC (Extended Data Fig. 1b), and from classical anatomical landmarks^22^. The proportion of the number of fibers counted in the nA vs. preBötC is represented (n = 6 for nA, n = 6 for preBötC, 3 sections counted per animal, Extended Data Fig. 1c, d). Scale bar, 50 µm. **b**, Retrograde tracing with Cholera Toxin B to localize OT neurons that project onto preBötC/nA neurons (n = 2, bilateral injections) (Extended Data Fig. 1e-g). Scale bars, large image 100 µm, inset 50 µm. **c**, ChR2-tdTomato expression in OT neurons, with very little, and likely overestimated, non-specific expression (Extended Data Fig. 2a). ChR2-tdTomato was detected in fibers in the preBötC/nA, enabling local photoexcitation. ChAT, choline acetyltransferase. Scale bars, left 100 µm, right 50 µm. **d**, Strategy for bilateral photoexcitation of OT fibers in the preBötC/nA. Optical fibers were positioned following electrophysiological mapping of the preBötC. NK1-R, neurokinin 1 receptor. Scale bar 500 µm. **e**, Photoexcitation (1 min, 50 Hz, 10 ms pulses) of OT fibers in the preBötC/nA of freely moving mice placed individually in plethysmography chambers to record respiratory activity, and previously implanted with ECG telemetry probes to measure HR. Adult OT::Cre;Ai27(LSL-ChR2) (Cre^+^, n = 8) and Ai27(LSL-ChR2) female littermates (Cre^-^, n = 4) were used. Gray shaded areas on the left trace correspond to the recordings shown in the middle trace. bpm, beats per minute. **f**, Photoexcitation of OT fibers in the preBötC/nA of Cre^+^ mice induced an amplification of RSA, a decrease in mHR, an increase in respiratory frequency (freq) and no change in respiratory amplitude (ampl). Each point corresponds to the average of ∼5 photoexcitation trials per mouse. RM one-way ANOVA, Tukey’s multiple comparison. cpm, cycles per minute. au, arbitrary units. **g**, Quantification of the relative effects (delta (*Δ*) in %) induced by photoexcitation of OT fibers in Cre^+^ (from absolute values shown in **f**) and Cre^-^ (absolute values not shown) mice. Cre^-^ mice showed no changes in any parameters measured. Intra-group delta (*Δ*) changes (photoexcitation *vs.* pre-photoexcitation, shown in **f**), RM one-way ANOVA, Tukey’s multiple comparison. Inter-group comparison (Cre^+^ *vs*. Cre^-^), unpaired t-test. **h**, The delta (*Δ*) increase in RSA amplitude upon OT fibers’ photoexcitation is correlated with the absolute RSA amplitude during the pre-photoexcitation control period, while the delta decrease in mHR is not correlated to the pre-photoexcitation mHR level. Pearson correlation analysis, simple linear regression plotted.

Next we examined whether these PVN^OT^ fibers in the preBötC/nA can modulate cardiorespiratory function, and in particular RSA. We used OT::Cre;Ai27(LSL-ChR2) adult female mice to optogenetically stimulate PVN^OT^ fibers in the preBötC/nA bilaterally, *in vivo* in freely moving condition (Fig. 1c-d, Extended Data Fig. 2a, b). Cardiorespiratory functions were assessed with telemetry ECG and whole-body plethysmography. Prolonged photoexcitation (1 min, 50 Hz, 10 ms pulses, 10 mW) in animals during rest induced a strong and sustained amplification of RSA (+56 %, from 13.2 ± 3.2 to 19.5 ± 4.3 bpm) and a sustained decrease in mHR (-7 %, from 532 ± 18 to 497 ± 19 bpm). However, it had only a moderate respiratory effect, inducing a small increase in respiratory frequency (+5 %, from 125 ± 9 to 130 ± 9 cpm) with no change in respiratory amplitude (Fig. 1e-g). There was no effect of photostimulation in control mice. These data show that PVN^OT^→preBötC/nA neurons can amplify RSA and decrease mHR in mice at rest.

### OT release in the preBötC/nA by PVN^OT^→preBötC/nA neurons induces the amplification of RSA

OT fibers can release OT, but also other neurotransmitters such as glutamate^23,24^. To determine if the cardiorespiratory effects evoked by photoexcitation of PVN^OT^ fibers are due specifically to a release of OT in the preBötC/nA, we used anesthetized OT::Cre;Ai27(LSL-ChR2) adult mice and recorded ECG and inspiratory EMG. Unilateral photoexcitation of PVN^OT^ fibers in the preBötC/nA induced a RSA amplification (+53 %, from 3.8 ± 0.5 to 5.8 ± 0.8 bpm in females; +50 %, from 6.1 ± 0.9 to 9.0 ± 1.3 bpm in males) and a mHR decrease (-2 %, from 269.5 ± 11.9 to 264.5 ± 12.04 bpm in females; -3 %, from 214.1 ± 6.6 to 207.8 ± 6.6 bpm in males) (Fig. 2a-c), of slightly reduced magnitudes compared to bilateral photoexcitation in conscious mice (Fig. 1e-g). There was no change in respiratory frequency, but a tendency for an increase in inspiratory amplitude. As female and male mice had responses of similar profiles and magnitudes (Fig. 2c, Extended Data Fig. 3a), mixed sexes were used in all cohorts throughout the rest of this study (unless specified). A selective OT receptor (OT-R) antagonist ((d(CH_2_)_5_^1^,Tyr(Me)^2^,Thr^4^,Orn^8^,des-Gly-NH_2_^9^)-Vasotocin)^23,25^ was then injected unilaterally in the preBötC/nA (Fig. 2i), which upon photoexcitation at the same site almost abolished the RSA amplification, while the decrease in mHR persisted (Fig. 2e-g, Extended Data Fig. 3b-d). Photoexcitation following injection of vehicle (Fig. 2e, g, Extended Data Fig. 3c-d and 4a-b), or photoexcitation of the preBötC/nA contralateral to the OT-R antagonist injection site (Fig. 2e-g, Extended Data Fig. 3b-d), induced similar cardiorespiratory effects as in the control pre-injection condition. These data show that PVN^OT^ neurons can induce RSA amplification by releasing OT in the preBötC/nA, whereas the mHR effect is dependent on a different mechanism.

**Fig. 2:**
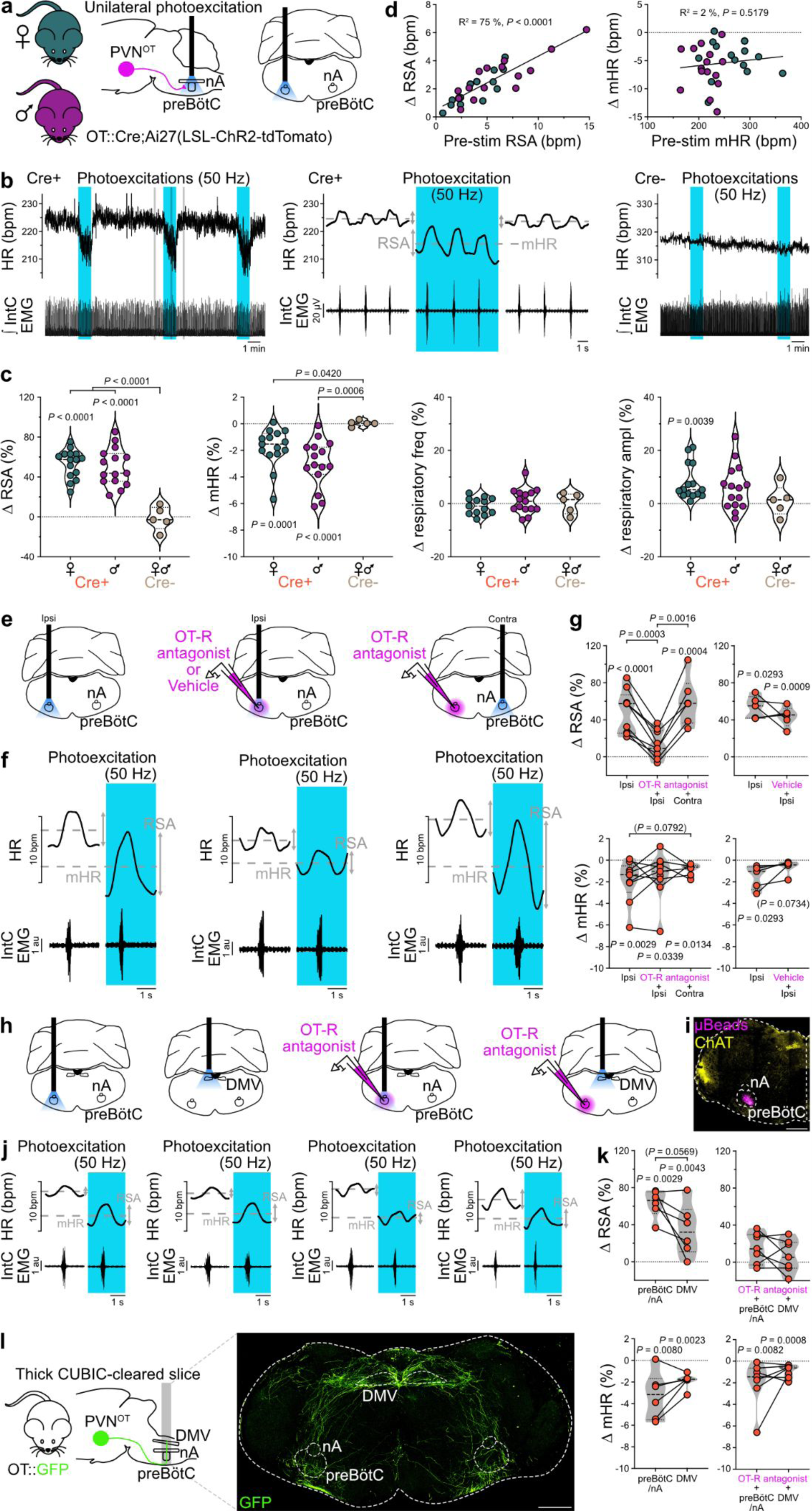
PVN^OT^ fibers can amplify RSA by releasing OT in the preBötC/nA, and can decrease mHR *via* DMV connectivity, in adult anesthetized mice. **a-c**, Unilateral photoexcitation of PVN^OT^ fibers in the preBötC/nA of anesthetized OT::Cre;Ai27(LSL-ChR2) adult female and male mice (**a**) induces RSA amplification and mHR decrease (**b**, **c**) (n = 15 Cre^+^ females, n = 15 Cre^+^ males, n = 5 Cre^-^ females and males). Intra-group delta (*Δ*) changes (photoexcitation *vs.* pre-photoexcitation), RM one-way ANOVA, Tukey’s multiple comparison. Inter-group comparison (Cre^+^ females *vs*. Cre^+^ males *vs*. Cre^-^), one-way ANOVA, Tukey’s multiple comparison, except for mHR and respiratory amplitude, Kruskal-Wallis test, Dunn’s multiple comparison. Raw data are shown in Extended Data Fig. 3a. IntC EMG, intercostal electromyography. **d**, The delta increase in RSA amplitude resulting from OT fibers’ photoexcitation is correlated with the absolute RSA amplitude during the pre-photoexcitation control period, while the delta decrease in mHR is not correlated to the pre-photoexcitation mHR level. Pearson correlation analysis, simple linear regression plotted. **e**-**g**, Ipsilateral (ipsi) injection of a selective OT receptor (OT-R) antagonist ((d(CH_2_)_5_^1^,Tyr(Me)^2^,Thr^4^,Orn^8^,des-Gly-NH_2_^9^)-Vasotocin, 200 nl at 1 µM) in the preBötC/nA (**e**) abolishes the photoexcitation-induced RSA amplification, but not the mHR decrease (**f**, **g**). Photoexcitation following ipsilateral vehicle injection (**g**, traces shown in Extended Data Fig. 4a), and photoexcitation in the contralateral (contra) preBötC/nA, induced similar RSA and mHR effects as ipsilateral pre-injection photoexcitation (n = 9 for “ipsi” and “OT-R antagonist + ipsi”, n = 6 for “ipsi” and “OT-R antagonist + ipsi” and “OT-R antagonist + contra”, n = 5 for “ipsi” and “vehicle + ipsi”). Intra-condition delta (*Δ*) changes (photoexcitation *vs.* pre-photoexcitation), RM one-way ANOVA, Tukey’s multiple comparison. Inter-conditions comparisons for OT-R antagonist (“ipsi” *vs.* “OT-R antagonist + ipsi” *vs.* “OT-R antagonist + contra”), RM mixed-effects analysis with the Geisser-Greenhouse correction, Tukey’s multiple comparison. Inter-conditions comparisons for vehicle (“ipsi” *vs.* “vehicle + ipsi”), paired t-test. Traces in **f** were obtained in the conditions shown above in **e**, following OT-R antagonist injection for the middle trace. Raw data for RSA and mHR, and data for the effects on respiratory frequency and respiratory amplitude, are shown in Extended Data Fig. 3b-d. Localisation of injection spots are shown in Extended Data Fig. 4b. **h**-**k**, Photoexcitation of PVN^OT^ fibers either in the preBötC/nA or in the DMV induce the same effects, both before or after injection of the OT-R antagonist in the preBötC/nA (n = 6 for “preBötC/nA” and “DMV” before OT-R antagonist, n = 7 for “preBötC/nA” and “DMV” after OT-R antagonist). Intra-condition delta (*Δ*) changes (photoexcitation *vs.* pre-photoexcitation), RM one-way ANOVA, Tukey’s multiple comparison. Inter-conditions comparisons before and after OT-R antagonist injection (“preBötC/nA” *vs.* “DMV”), paired t-test. Traces in **j** were obtained in the conditions shown above in **h**. Raw data for RSA and mHR, and data for the effects on respiratory frequency and respiratory amplitude, are shown in Extended Data Fig. 3e-g. **i**, Fluorescent microbeads in the injectate solutions enabled the localisation of the injection spots for the OT-R antagonist or vehicle *a posteriori*. Scale bar, 500 µm. **l**, CUBIC clearing of thick (1 mm) coronal brainstem slices at the level of the preBötC/nA in OT::GFP mice (n = 3) shows OT fibers crossing both the preBötC/nA and the DMV. Scale bar, 500 µm.

Interestingly, the larger the RSA amplitude was prior to the photoexcitation of PVN^OT^ fibers in the preBötC/nA, the larger the delta in RSA amplification was during photoexcitation, with a strong correlation between the two parameters occurring in both the freely moving and anesthetized conditions (Fig. 1h and 2d). This suggests that OT likely acts in the preBötC/nA as a RSA gain amplifier. By contrast, the photoexcitation-induced decrease in mHR was not correlated with the pre-photoexcitation mHR level, further pointing towards a different underlying mechanism. OT was previously shown to act in the DMV to decrease mHR^26^. Photoexcitation of OT fibers in the DMV before and after the injection of the OT-R antagonist in the preBötC/nA induced the same effects as the preBötC/nA photoexcitations, with RSA amplification only before the injection of the OT-R antagonist, and mHR decrease both before and after the antagonist injection (Fig. 2h-k, Extended Data Fig. 3e-g). This suggests that some OT fibers could contact both the preBötC/nA and the DMV to amplify RSA and decrease mHR, respectively. This is supported at the anatomical level, as tracing of OT fibers in CUBIC-cleared thick brainstem slices of OT::GFP mice containing the preBötC/nA and the DMV showed OT fibers crossing both structures (Fig. 2l).

### In the preBötC/nA, a subgroup of inhibitory preBötC neurons expresses the OT-R

So far, our data show that PVN^OT^ fibers can amplify RSA by releasing OT in the preBötC/nA, most likely acting on preBötC rather than nA neurons because most PVN^OT^ projection fibers were found in the preBötC (Fig. 1a, Extended Data Fig. 1c-d). To identify the cells modulated by OT to induce RSA amplification, we used OT-R::Cre;Ai14(LSL-tdTomato) adult mice to label OT-R^+^ cells (Fig. 3a-e, Extended Data Fig. 5a-d). In the preBötC/nA, very few OT-R^+^ neurons are nA neurons (7.6 ± 3.8 neurons on each side) or nA^Cardiac^ neurons (2.0 ± 0.9 neurons on each side), identified as nA neurons not expressing the calcitonin gene-related peptide^27,28^ - by contrast, most DMV neurons are OT-R^+^ (587.2.8 ± 106.4 neurons on each side) (Fig. 3a, Extended Data Fig. 5a). OT-R^+^ cells in the preBötC/nA are predominantly neurons (66.0 ± 4.3 % neurons vs. 12.1 ± 2.6 % astrocytes), with qualitatively small somata (Fig. 3b, Extended Data Fig. 5b). They are present predominantly within the anatomical boundaries of the preBötC, expressing classical markers of the preBötC neuronal population, including the expression of neurokinin 1 and µ-opioid receptors (Fig. 3b, Extended Data Fig. 5c). However, few contralaterally-projecting preBötC neurons were OT-R^+^. PreBötC^OT-R^ neurons represent a population of ∼700 neurons bilaterally (357.5 ± 4.4 neurons on each side). They are localized throughout the preBötC, with no specific clustering, but appear as a specific population within the respiratory cell column since few OT-R^+^ neurons were found in the adjacent Bötzinger Complex (Fig. 3c). These data show that the OT-induced amplification of RSA is most likely mediated via PVN^OT^→preBötC^OT-R^ connectivity.

**Fig. 3:**
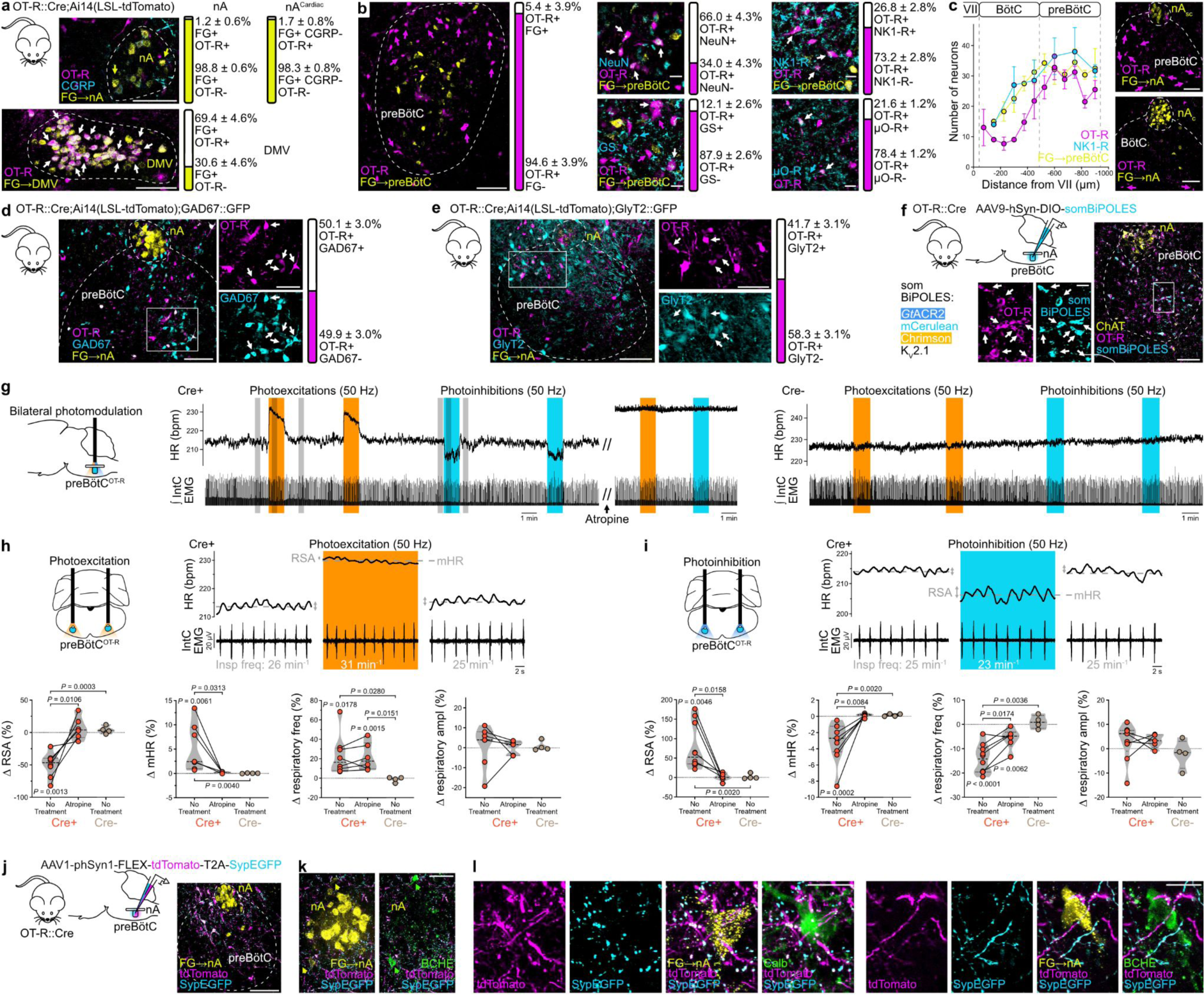
Anatomo-functional characterisation of OT-R^+^ cells in the preBötC/nA controlling RSA amplitude. **a**, Crossing of OT-R::Cre mice with Ai14(LSL-tdTomato) mice to express tdTomato only in OT-R^+^ cells. FG was injected intraperitoneally, to label nA and DMV neurons (n = 6 mice for nA counts, n = 5 mice for DMV counts). nA^Cardiac^ neurons were detected as nA FG^+^ neurons that do not express the calcitonin gene-related peptide (CGRP^-^). Arrows indicate labeled neurons, with white arrows showing double labeled OT-R^+^ FG^+^ neurons. Individual data are shown in Extended Data Fig. 5a. Scale bars, 100 µm. **b**, Unilateral preBötC boundaries were mapped using contralateral preBötC FG injection (Extended Data Fig. 1b and c), and using classical anatomical landmarks of the preBötC location^22^. PreBötC^OT-R^ (OT-R^+^) cells were tested for their contralateral projection phenotype (FG^+^, n = 4 mice), neuronal vs. astrocytic phenotype (NeuN^+^ vs. Glutamine Synthetase (GS)^+^, respectively, n = 4 mice), and for the expression of two classical markers of preBötC neurons (NK1-R^+^, n = 4 mice; µO-R^+^, n = 6 mice). Arrows indicate labeled neurons, with white arrows showing double labeling between OT-R^+^ and the cyan marker. Individual data are shown in Extended Data Fig. 5b and c. Scale bar for preBötC^OT-R^ and FG only, 100 µm; other scale bars, 20 µm. **c**, OT-R^+^ cells are densely present in the preBötC, and little present in the adjacent Bötzinger Complex (BötC). nA_sc_, semi-compact formation of the nA; nA_c_, compact formation of the nA. Scale bars, 100 µm. **d-e**, Crossing of OT-R::Cre;Ai14(LSL-tdTomato) mice with GAD67::GFP mice (**d**) or GlyT2::GFP mice (**e**) to label OT-R^+^ cells (tdTomato^+^) and GABAergic or glycinergic neurons (GFP^+^), respectively (n = 4 mice for GABAergic counts, n = 4 mice for glycinergic counts). Individual data are shown in Extended Data Fig. 5d. Scale bars full images, 100 µm; scale bars insets, 50 µm. **f**, Viral-mediated expression of the somBiPOLES optogenetic construct in preBötC^OT-R^ neurons of OT-R::Cre mice. Viral injections in Cre^-^ mice did not induce somBiPOLES expression (Extended Data Fig. 5e). An anti-OT-R antibody was used for immunohistochemical labeling. Scale bar full image, 100 µm; scale bar inset, 20 µm. **g**, Effects of bilateral photomodulations of preBötC^OT-R^ neurons in an anesthetized mouse (Cre^+^) on HR and inspiratory activity (IntC EMG), before and after intraperitoneal injection of the muscarinic receptor antagonist atropine (10 mg/kg, 600 µl), and compared to a control (Cre^-^) mouse. **h-i**, Expanded traces from the gray shaded areas in (**g**), and individual data showing bidirectional cardiorespiratory effects of photoexcitation (**h**) vs. photoinhibition (**i**) of preBötC^OT-R^ neurons (n = 8 Cre^+^ mice for photoexcitation and n = 10 Cre^+^ mice for photoinhibition without treatment, n = 6 Cre^+^ mice for photoexcitation/photoinhibition before vs. after atropine, n = 4 Cre^-^ mice). Intra-group delta (*Δ*) changes (photoexcitation *vs.* pre-photoexcitation), RM one-way ANOVA, Tukey’s multiple comparison, except for Cre^-^ RSA photoinhibition, Friedman test, Dunn’s multiple comparison. Cre^+^ before and after atropine comparison, paired t-test, except for respiratory amplitude photoexcitation, Wilcoxon matched-pairs signed rank test. Inter-group comparison (Cre^+^ no treatment *vs*. Cre^+^ atropine *vs*. Cre^-^ no treatment), Kruskal-Wallis test, Dunn’s multiple comparison. Raw data are shown in Extended Data Fig. 5f and g. **j**, Viral-mediated expression of tdTomato and synaptophysin-enhancedGFP (SypEGFP) in preBötC^OT-R^ neurons. Scale bar, 100µm. **k-l**, PreBötC^OT-R^ neurons project to (tdTomato^+^ fibers) and make putative pre-synaptic contacts with (SypEGFP^+^ puncta) nA^Cardiac^ neurons (FG^+^ BCHE^+^ and FG^+^ Calb^+^ neurons). BCHE, butyrylcholinesterase; Calb, calbindin. Scale bar in (**k**), 50 µm; scale bars in (**l**), 20 µm.

A cumulative body of work suggests that the inspiratory component of RSA is generated by inhibitory preBötC neurons^10–12,14,29^. Also, OT was shown to act selectively on inhibitory neurons in neuronal networks of the hippocampus and the amygdala, to amplify the gain of inhibitory synaptic connectivity^25,30,31^. Accordingly, we tested whether preBötC^OT-R^ neurons express the two classical inhibitory neurotransmitters GABA or glycine, using OT-R::Cre;Ai14(LSL-tdTomato);GAD67::GFP mice, OT-R::Cre;Ai14(LSL-tdTomato);GlyT2::eGFP mice and immunolabelling (Fig. 3d-e). We found in adult mice that 50% of preBötC^OT-R^ neurons are GABAergic (Fig. 3d, Extended Data Fig. 5d), and 42% are glycinergic (Fig. 3e, Extended Data Fig. 5d). Therefore, preBötC^OT-R^ neurons are predominantly inhibitory.

### PreBötC^OT-R^ neurons form a cardiorespiratory hub that controls RSA amplitude *via* preBötC^OT-R^→nA^Cardiac^ projections

Our data show that PVN^OT^ neurons can amplify RSA by releasing OT onto preBötC^OT-R^ neurons, which are predominantly inhibitory, and not by directly modulating the output nA^Cardiac^ neurons. PreBötC neurons are defined as a group of neurons generating the inspiratory rhythm^18–20^. To determine the functional role of preBötC^OT-R^ neurons for cardiorespiratory activities, we injected a Cre-dependent adeno-associated virus (AAV) bilaterally in the preBötC of adult OT-R::Cre mice, to express the somBiPOLES construct that enables bidirectional optogenetic modulation^32^ (Fig. 3f, Extended Data Fig. 5e). Prolonged photoexcitation of preBötC^OT-R^ neurons in anesthetized mice induced a rapid and sustained decrease in RSA amplitude (-49.9 ± 6.6 %), an increase in mHR (+5.3 ± 1.8 %) and an increase in respiratory frequency (+23.2 ± 7.3 %) (Fig. 3g-h, Extended Data Fig. 5f). Prolonged photoinhibition of the same neurons induced the opposite effects, causing a rapid and sustained increase in RSA amplitude (+77.2 ± 18.5 %), a decrease in mHR (-3.1 ± 0.7 %) and a decrease in respiratory frequency (-14.3 ± 1.8 %) (Fig. 3g and i, Extended Data Fig. 5g). There was no effect on respiratory amplitude, nor on all parameters in control mice. Interestingly, these bidirectional effects on RSA amplitude indicate that preBötC^OT-R^ neurons continuously set an ongoing level of RSA amplitude. The large variations in the amplitude of these bidirectional effects indicates that external modulators like OT can induce strong effects on RSA amplitude by acting on preBötC^OT-R^ neurons. Moreover, preBötC^OT-R^ neurons are not part of the preBötC inspiratory rhythmogenic subgroup, since the photoinhibition of preBötC^OT-R^ neurons did not terminate respiratory activity, in contrast to an inactivation of preBötC rhythmogenic neurons^12,33,34^. Therefore, preBötC^OT-R^ neurons form a core cardiorespiratory hub within the preBötC that controls RSA amplitude.

Intraperitoneal injection of the muscarinic receptor antagonist atropine, which blocks parasympathetic activity to the heart, blocked the effects of optogenetic modulations of preBötC^OT-R^ neurons on RSA and mHR, but not the respiratory effects (Fig. 3g-i, Extended Data Fig. 5f and g). This shows that preBötC^OT-R^ neurons modulate cardiac activity *via* modulation of cardiac parasympathetic activity, therefore most likely *via* nA^Cardiac^ neurons because they are the sole cardiac parasympathetic preganglionic neurons that are respiratory modulated^9–11,14^. Also, the predominantly inhibitory nature of preBötC^OT-R^ neurons (glycinergic and GABAergic) (Fig. 3d and e, Extended Data Fig. 5d), in addition to the direction of mHR effects during the photomodulation of preBötC^OT-R^ neurons (increased mHR during photoexcitation, decreased mHR during photoinhibition) (Fig. 3g-i, Extended Data Fig. 5f and g), strongly suggest that the inputs from preBötC^OT-R^ neurons to cardiac parasympathetic preganglionic neurons are monosynaptic. To determine if preBötC^OT-R^ neurons project onto nA^Cardiac^ neurons, we injected a Cre-dependent AAV in OT-R::Cre mice to express tdTomato and synaptophysin-GFP in preBötC^OT-R^ neurons, and we analyzed the preBötC^OT-R^→nA^Cardiac^ axonal projections (tdTomato^+^) and putative pre-synaptic contacts (synaptophysin-GFP^+^) (Fig. 3j-l). To identify specifically the nA^Cardiac^ neurons, we used intraperitoneal FluoroGold injection to label all nA neurons, and selective immunolabelling of nA^Cardiac^ neurons based on the recent identification of two specific markers, calbindin and butyrylcholinesterase^28^ (Fig. 3k and l), and the confirmation of a third marker with the absence of expression of the calcitonin gene-related peptide^27,28^. We found dense preBötC^OT-R^ axonal projections throughout the rostrocaudal extent of the nA. All identified nA^Cardiac^ neurons (51 butyrylcholinesterase^+^ and 17 calbindin^+^ nA^Cardiac^ neurons bilaterally in 4 mice) received projections and putative pre-synaptic contacts from preBötC^OT-R^ neurons (Fig. 3l). Therefore, these data are consistent with a direct monosynaptic connectivity from preBötC^OT-R^ neurons to nA^Cardiac^ neurons.

### The preBötC^OT-R^→nA^Cardiac^ inspiratory connectivity is glycinergic and amplified by OT

Our results thus far show that preBötC^OT-R^ neurons are predominantly inhibitory, make putative monosynaptic contacts with nA^Cardiac^ neurons, and regulate cardiorespiratory functions and RSA amplitude. It is therefore likely that the amplification of RSA by OT, identified by our optogenetic stimulations of PVN^OT^→preBötC fibers, is due to an amplification of the preBötC^OT-R^→nA^Cardiac^ inspiratory and inhibitory connectivity. To test this possibility, we used brainstem slices from newborn mice that contain the preBötC/nA structures, and in which preBötC neurons continue spontaneously to generate rhythmic inspiratory bursting activity^18,35,36^. We first performed extracellular recordings of preBötC population activity, simultaneously with whole-cell current-clamp recordings of inspiratory bursting preBötC neurons (Fig. 4a-b). Bath application of the selective OT-R agonist [Thr^4^,Gly^7^]-OT (TGOT)^25,37,38^ induced an increase in preBötC inspiratory bursting frequency (+47.0 ± 20.4 %), along with an increase in preBötC network excitability as shown by the appearance of spiking activity between ongoing inspiratory bursts (Fig. 4a-c). To confirm that these effects are mediated specifically by TGOT’s action on preBötC^OT-R^ neurons, we used isolated preBötC “island” preparations^39^, in which TGOT still increased preBötC inspiratory bursting frequency (+40.1 ± 11.5 %) (Fig. 4d-e). Immunolabelling of preBötC^OT-R^ neurons in newborn OT-R::Cre;Ai14(LSL-tdTomato);GAD67::GFP and OT-R::Cre;Ai14(LSL-tdTomato);GlyT2::eGFP mice displayed a similar expression profile compared to adult mice, with a predominantly inhibitory phenotype (GABAergic and/or glycinergic) (Fig. 4f-g).

**Fig. 4:**
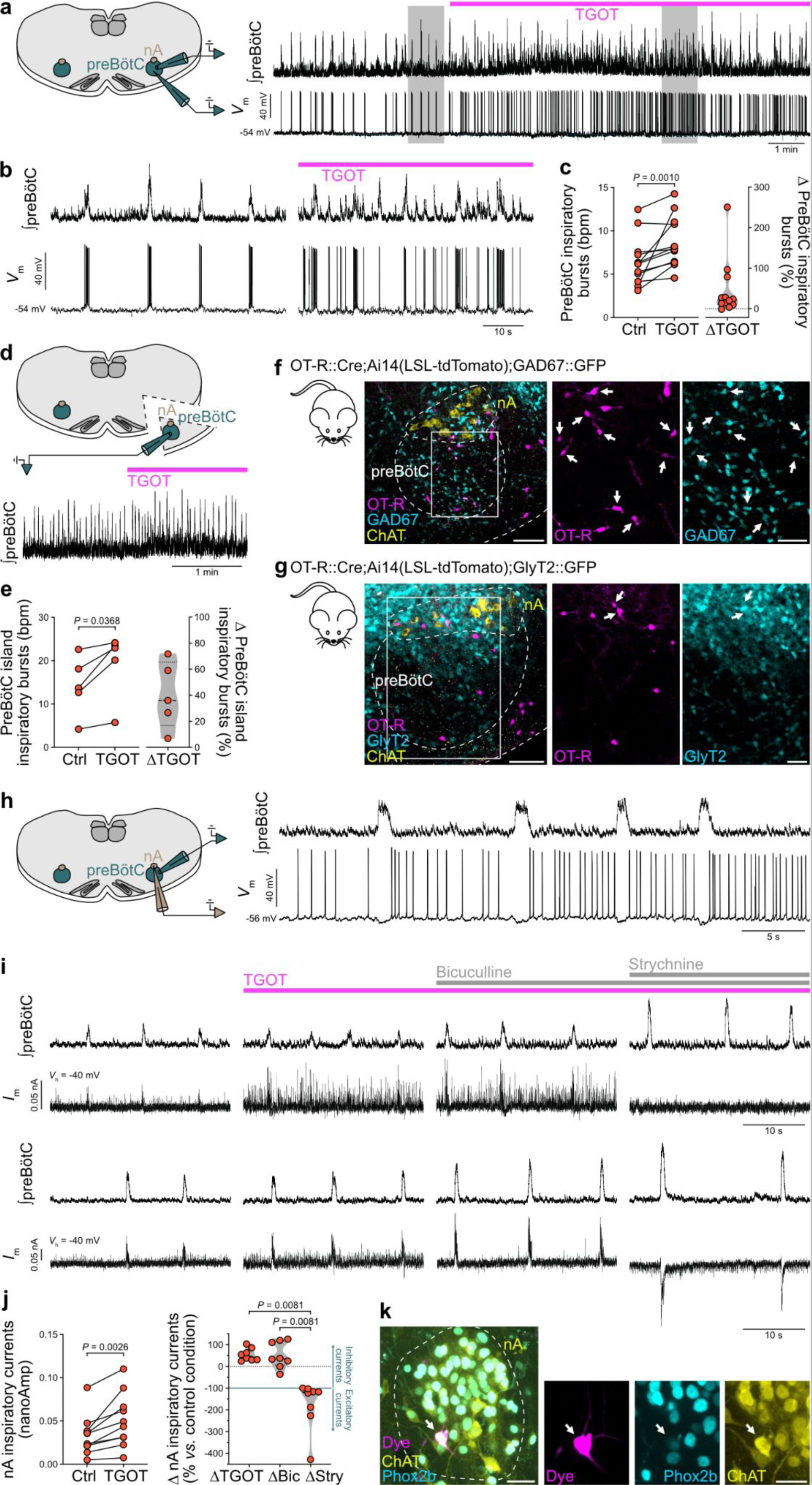
TGOT increases the frequency of preBötC inspiratory bursts and amplifies the inspiratory glycinergic currents occurring in nA neurons in rhythmic slices *in vitro*. **a**, Effects induced by the application of the selective OT-R agonist TGOT (0.5 µM) on preBötC population activity (⎰preBötC, extracellular recordings) and on the activity of an individual inspiratory neuron (*V*_m_, whole-cell current-clamp recordings), in a rhythmic preBötC slice from a neonatal mouse (all recordings performed at P0-P5). **b**, Expanded traces from the gray shaded areas in (**a**). **c**, TGOT increases the frequency of preBötC inspiratory bursts in rhythmic preBötC slices (n = 12 mice). Wilcoxon matched-pairs signed rank test. bpm, bursts per minute. **d**, Effects induced by the application of TGOT (0.5 µM) on preBötC population activity in a preBötC “island” preparation, where the preBötC is isolated from the rest of the slice. **e**, TGOT increases the frequency of preBötC inspiratory bursts in preBötC “island” preparations (n = 5 mice). Paired t-test. **f-g**, In P0 triple transgenic mice, preBötC^OT-R^ neurons show the same predominantly GABAergic (**f**) or glycinergic (**g**) phenotype as in adult mice (Fig. 3d-e) (arrows indicate OT-R and GAD67 or GlyT2 co-labelling). Scale bars full images, 100 µm; scale bars insets, 50 µm. **h**, Simultaneous recordings of preBötC population activity and of the activity of a typical respiratory-modulated nA neuron (*V*_m_, whole-cell current-clamp recordings). All respiratory-modulated nA neurons (n = 10 mice) were hyperpolarized during the inspiratory preBötC bursts, causing inhibition of their spiking activity. **i**, Successive effects induced by the application of TGOT (0.5 µM), bicuculline (GABA_A_ receptor antagonist, 10 µM) and strychnine (glycine receptors antagonist, 5 µM) on preBötC population activities and on the currents recorded from respiratory-modulated nA neurons (*I*_m_, whole-cell voltage-clamp recordings; *V*_h_, -40 mV holding voltage). The top and bottom traces are representative examples of the two types of effects found following strychnine application. **j**, All recorded respiratory-modulated nA neurons showed volleys of inhibitory currents during preBötC inspiratory bursts in the control condition, which were amplified by TGOT application (n = 10 mice). During TGOT application, bicuculline application did change the delta (*Δ*) amplification of the inspiratory inhibitory currents relative to control condition (n = 8 mice), while strychnine application either abolished all currents (n = 5 mice), or exposed the presence of volleys of excitatory currents during preBötC inspiratory bursts (n = 3 mice). Control vs. TGOT, paired t-test. Delta changes following TGOT, bicuculline and strychnine applications, Friedman test, Dunn’s multiple comparison. **k**, Example of a recorded nA neuron (arrows indicate the dye^+^ recorded neuron that is Phox2b^+^ and ChAT^+^). Scale bar full image, 40 µm; scale bar inset, 20 µm.

Then, to assess the effect of TGOT application on the preBötC^OT-R^→nA^Cardiac^ connectivity, we performed simultaneous extracellular recordings of preBötC population activity, and whole-cell current- and voltage-clamp recordings of neurons within the nA boundaries that presented inspiratory modulation of their activity (Fig. 4h-i). *Post hoc* immunolabelling confirmed that all recorded neurons were nA neurons (Fig. 4k). As shown previously, inspiratory-modulated nA^Cardiac^ neurons display spiking activity during the expiratory phase of the respiratory cycle, which is silenced during inspiratory bursts by hyperpolarizing synaptic currents^10,11^. We recorded from nA neurons that were all active during the expiratory period and hyperpolarized during preBötC inspiratory bursts (current-clamp), due to large outward inhibitory currents (voltage-clamp, -40 mV holding voltage) (Fig. 4h-i). TGOT application amplified the volleys of inspiratory inhibitory currents in all recorded nA neurons (+60.6 ± 11.1 %, n = 10 neurons, each from a different mouse) (Fig. 4i-j). Application of the GABA_A_ receptor antagonist bicuculline had no effect, while application of the glycine receptors antagonist strychnine abolished the inspiratory inhibitory currents in all nA neurons. Following strychnine application, half of these nA neurons showed an absence of remaining currents, while the other half showed excitatory currents during preBötC inspiratory bursts that were undetectable before strychnine application (blocked by the AMPA/kainate receptor antagonist CNQX, data not shown). Also, a post-inhibitory rebound was evident after each volley of inspiratory inhibitory currents on nA neurons (Fig. 4i), inducing an increase in spiking activity immediately after each inspiratory burst (Fig. 4h), which tended to be amplified by TGOT. Together, these electrophysiological data show that OT raises the excitability of inspiratory preBötC neurons, leading to the amplification of the preBötC→nA^Cardiac^ inspiratory glycinergic connectivity.

### The OT-induced RSA amplification is mediated *via* amplified respiratory modulation of cardiac parasympathetic activity from nA^Cardiac^ neurons to the heart

So far, our data show that a PVN^OT^→preBötC^OT-R;Glycine^→nA^Cardiac^ neuronal network can amplify RSA, with PVN^OT^ fibers releasing OT onto inhibitory preBötC^OT-R^ neurons to amplify the strength of the preBötC^OT-R^→nA^Cardiac^ inspiratory glycinergic connectivity. This strongly suggests that RSA is then amplified *via* an amplification of the inspiratory modulation of parasympathetic activity from nA^Cardiac^ neurons to the heart. However, preBötC neurons also provide inspiratory modulation of pre-sympathetic neurons of the rostral ventrolateral medulla oblongata, producing a respiratory modulation of blood pressure ^12,40,41^, which could indirectly regulate RSA amplitude. To address this possibility, we used the *in situ* Working Heart-Brainstem Preparation, which enables recording autonomic and respiratory nerve activities in a perfused and decerebrate preparation devoid of the depressant effect of anesthesia^13^. We employed rats because this preparation functions considerably better in rats than in mice, and because the difficult dissection and recording of the vagal branch that provides parasympathetic activity to the heart is more achievable in rats^42^. However, since all our work so far was done in mice, we first used *in vivo* anesthetized rats to confirm the reproducibility of results between both species. Injection of TGOT bilaterally in the preBötC of anesthetized rats induced an amplification of RSA (+52.7 ± 15.8 %), a small yet significant mHR decrease (-1.0 ± 0.4 %), no alteration in respiratory frequency and an increase in inspiratory amplitude (+7.0 ± 4.1 %) (Fig. 5a-c, Extended Data Fig. 6a-c). These effects are therefore very similar to those produced in anesthetized mice by OT release in the preBötC during optogenetic stimulation of PVN^OT^ fibers (Fig. 2). Then, using the *in situ* Working Heart-Brainstem Preparation, injection of TGOT bilaterally in the preBötC induced an amplification of RSA (+107.1 ± 17.8 %), no change in mHR, and no alteration in the frequency and amplitude of inspiratory and post-inspiratory discharges recorded from the phrenic and cervical vagus nerves, respectively (Fig. 5d-f, Extended Data Fig. 6d-f). TGOT injection in the preBötC induced no change in thoracic (T8-T10) sympathetic chain recordings, which carry vasomotor activity, and no change in perfusion pressure, which in this preparation is the equivalent to blood pressure. Conversely, TGOT injection in the preBötC induced an amplification of the respiratory modulation of cardiac parasympathetic activity (+73.6 ± 10.1 %), recorded from the left thoracic cardiac vagal branch^42^ (Fig. 5h-i, Extended Data Fig. 6g). These data show that OT in the preBötC induces RSA amplification *via* amplified respiratory modulation of cardiac parasympathetic activity to the heart, not *via* a modulation of sympathetic vasomotor activity.

**Fig. 5:**
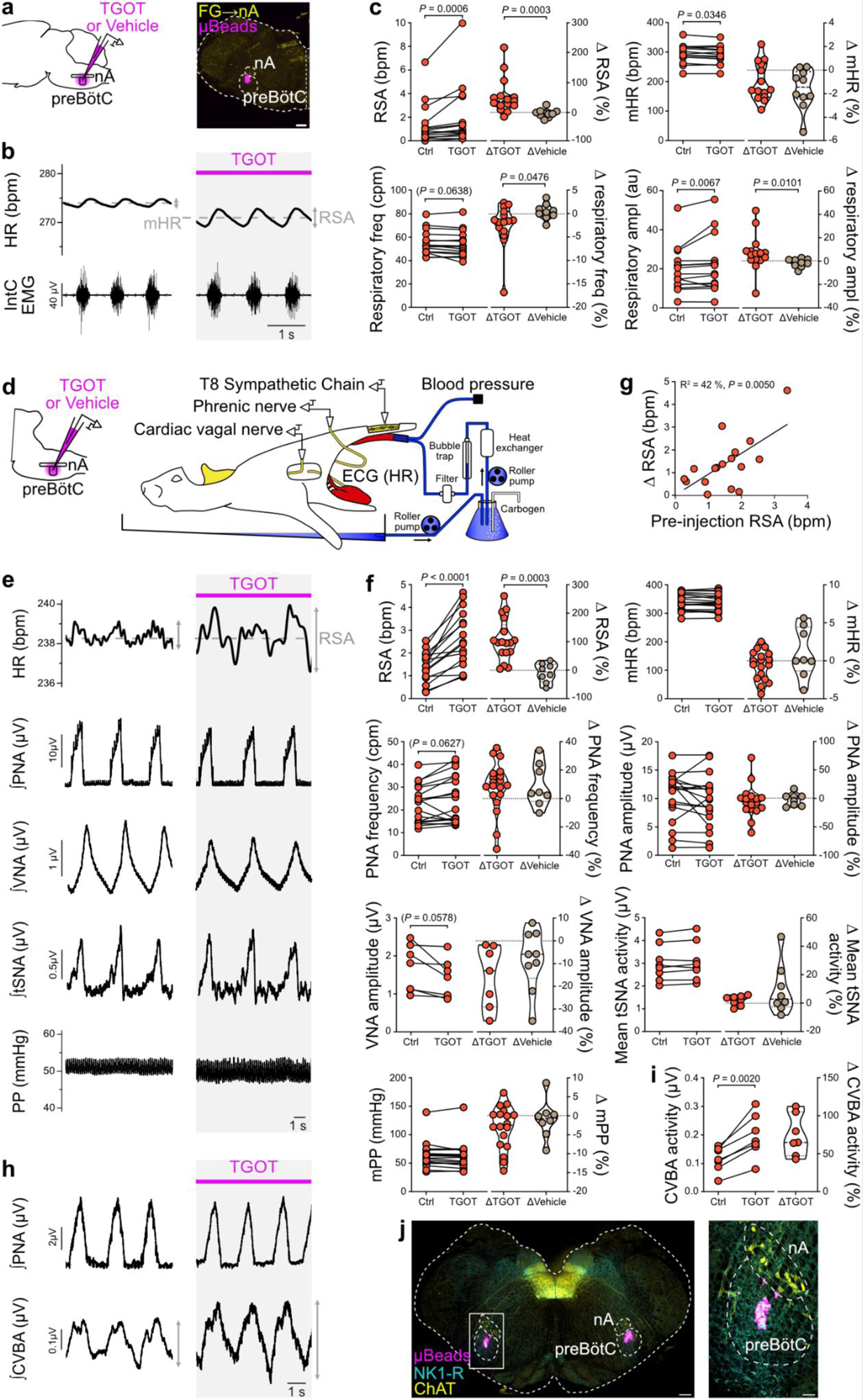
TGOT injection in the preBötC induces RSA amplification due to amplified respiratory modulation of cardiac parasympathetic activity in adult anesthetized rats and in *in situ* Working Heart-Brainstem Preparations of juvenile rats. **a**, Bilateral injection of TGOT (50 nl at 0.5 µM) or vehicle in the preBötC of adult anesthetized Wistar rats. Fluorescent microbeads in the injectate solutions enabled the localisation of the injection spots *a posteriori* (mapped in Extended Data Fig. 6c). Scale bar, 300 µm. **b-c**, The bilateral injection of TGOT in the preBötC induced an amplification of RSA, a decrease in mHR, an increase in respiratory amplitude (ampl) but no change in respiratory frequency (freq). Absolute values for the conditions before and after TGOT injection (n = 15), paired t-test for respiratory frequency and Wilcoxon matched-pairs signed rank test for RSA, mHR and respiratory amplitude. Relative effects (delta (*Δ*) in %) induced by TGOT or vehicle (n = 10, absolute values in Extended Data Fig. 6b) injections, Mann-Whitney test or unpaired t-test (mHR). IntC EMG, intercostal electromyography. **d**, Bilateral injection of TGOT or vehicle in the preBötC of *in situ* Working Heart-Brainstem Preparations of juvenile Wistar rats (P21-30). ECG, Electrocardiogram. **e-i**, Bilateral TGOT injection induced an amplification of RSA (**e**, **f**) and an increase in the respiratory modulation of the cardiac vagal branch activity (CVBA) (**h**, **i**), with no change in mHR, amplitude or frequency of the phrenic nerve activity (PNA), amplitude of the cervical vagal nerve activity (VNA), mean thoracic sympathetic nerve activity (tSNA) or mean perfusion pressure (PP). PNA, HR and PP were recorded in all *in situ* Working Heart-Brainstem Preparations, while VNA and tSNA (**e**, **f**), and CVBA (**h**, **i**), were recorded in different preparations. Integrated traces (**e**, **h**) are respiratory triggered averaging over >15 respiratory cycles. Absolute values for the conditions before and after TGOT injection (**f**, **i**) (RSA, n = 16; mHR, n = 18; PNA frequency, n = 19; PNA amplitude, n = 18; VNA, n = 7; tSNA, n = 8; mPP, n = 18; CVBA, n = 7), paired t-test or Wilcoxon matched-pairs signed rank test. Relative effects (delta (*Δ*) in %) induced by TGOT or vehicle injections (absolute values for vehicle injections in Extended Data Fig. 6e; RSA, n = 8; mHR, n=8; PNA frequency, n = 9; PNA amplitude, n = 8; VNA, n = 9; tSNA, n = 8; mPP, n = 8), Mann-Whitney test or unpaired t-test. The delta (*Δ*) increase in RSA amplitude following TGOT injection is correlated with the absolute RSA amplitude during the pre-injection control period (**g**). Pearson correlation analysis, simple linear regression plotted. **j**, Fluorescent microbeads in the injectate solutions enabled the localisation of the injection spots of TGOT or vehicle *a posteriori* (mapped in Extended Data Fig. 6h). NK1-R, neurokinin 1 receptor. ChAT, choline acetyltransferase. Scale bar large image 300 µm, inset 100 µm.

### OT neurons participate in the recovery of RSA amplitude during calming behavior following a stressful event

We have identified a multisynaptic neuronal network for OTergic amplification of RSA, from the hypothalamus to the brainstem to the heart. This network can be activated and induce RSA amplification during rest (Fig. 1e-g). Since RSA amplitude is increased during relaxation and calming behaviors^1,3^, we tested whether OT neurons can be activated endogenously to participate in the recovery of RSA amplitude during recovery from stress. We used OT::Cre;R26-LSL-hM4Di-DREADD male adult mice to chemogenetically inhibit OT neurons *in vivo* in freely moving condition (Fig. 6, Extended Data Fig. 7). Cardiovascular and respiratory functions were assessed with blood pressure telemetry recordings and whole-body plethysmography. Each mouse was habituated to a dedicated plethysmography chamber for a week, with nesting material from their home cage. At the end of this habituation period, the mice exhibited a predominantly calm and resting behavior in their plethysmography chamber. Then, the mice were injected intraperitoneally with either the DREADD agonist Compound 21 (C21), or vehicle, followed by a resting phase in their plethysmography chamber, a restraint stress test phase in a ventilated cylinder, and a recovery phase back in their plethysmography chamber (Fig. 6c). Compared to vehicle, C21 injections induced no change in RSA amplitude, blood pressure, or respiratory frequency and amplitude during the pre-stress resting phase (Fig. 6d-e, Extended Data Fig. 7). This shows that OT neurons do not influence RSA amplitude in resting condition. However, C21 injection decreased mHR compared to vehicle in both Cre^+^ (-8.2 ± 2.3 %) and Cre^-^ (-6.6 ± 0.9 %) mice, indicating an off-target effect of C21 on mHR. While all DREADD agonists have been shown to exhibit off-target effects^43^, importantly here RSA amplitude and respiratory parameters were not affected.

**Fig. 6:**
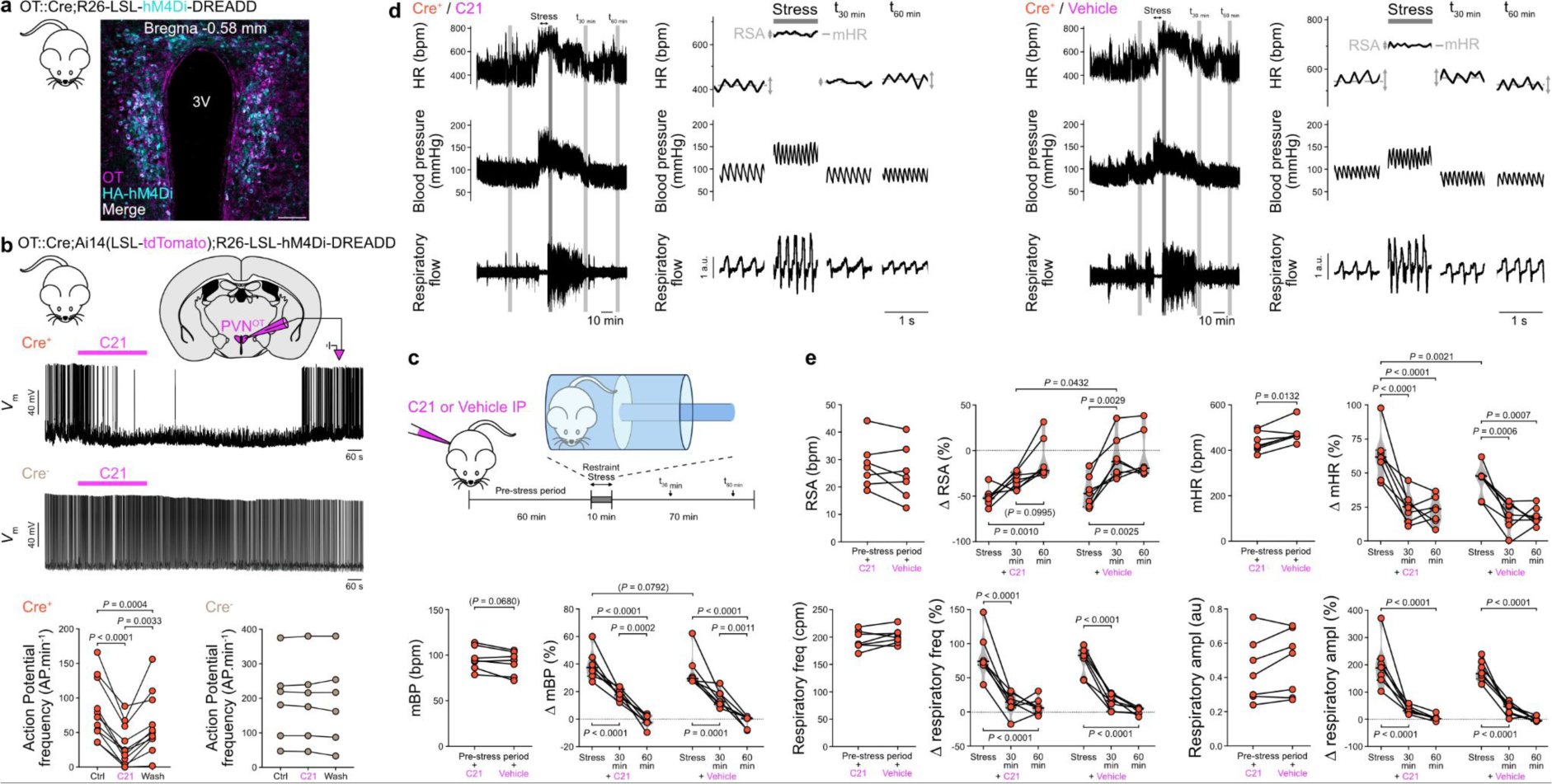
Chemogenetic inhibition of OT neurons slows down the recovery of RSA amplification following a restraint stress test. **a**, Expression of the HA-tagged inhibitory chemogenetic actuator hM4Di in OT neurons in adult transgenic mice. hM4Di is a G_αi/o_-protein coupled receptor from the Designer Receptors Exclusively Activated by Designer Drugs (DREADD) system. Scale bar, 100 µm. **b**, Whole-cell current-clamp recordings of PVN^OT^ neurons expressing tdTomato and hM4Di in slices from Cre^+^ mice, showing that application of the DREADD agonist Compound 21 (C21, 10 µM, for 5 min) inhibits action potentials. C21 had no effect on neurons recorded in the PVN in Cre^-^ mice (n = 3 Cre^+^ and n = 2 Cre^-^ P20 mice, at least 3 recorded neurons per mouse, each from a different slice). RM one-way ANOVA with the Geisser-Greenhouse correction, Tukey’s multiple comparison. **c**, Experimental strategy to test the effect of chemogenetic inhibition of OT neurons on respiratory and cardiovascular parameters, before and after a restraint stress test in a ventilated cylinder. The mice were implanted with blood pressure telemetry probes to enable measurements of blood pressure and HR. They were placed in a plethysmography chamber (to which they were habituated for 1 week) during the pre-stress and post-stress conditions, to enable measurement of their respiratory activity. **d-e**, Effects induced by the intraperitoneal injection of the hM4Di agonist Compound 21 (C21) or vehicle in adult Cre^+^ mice (n = 7, three trials averaged for each condition, C21 and vehicle, in each mouse; C21 or vehicle trials spaced by at least 24h). Traces on the right for both the C21 and vehicle conditions are enlargements of the grayed areas on their respective traces on the left (**d**). Chemogenetic inhibition of OT neurons did not alter RSA amplitude at rest, but slowed down the recovery of RSA amplification following the restraint test. Pre-stress conditions, C21 *vs.* vehicle, paired t-test (**e**). Stress and post-stress delta (*Δ* in %) effects compared to the pre-stress condition, C21 *vs.* vehicle, RM two-way ANOVA, Sidak’s multiple comparison. Cre^-^ data are shown in Extended Data Fig. 7.

Restraint stress induced a strong decrease (by ∼50%) in RSA amplitude of the same magnitude in all groups, and a strong increase in mHR, blood pressure and respiratory parameters (Fig. 6c-e, Extended Data Fig. 7). During the recovery from stress, chemogenetic inhibition of OT neurons specifically delayed the recovery in RSA amplitude, but not the return to basal values of other cardiovascular and respiratory parameters. Indeed, in vehicle-injected Cre^+^ mice, and in all Cre^-^ mice, RSA amplitude, mHR, respiratory frequency and amplitude had returned to baseline values 30 min following the stress test. In C21-injected Cre^+^ mice, all parameters followed the same recovery kinetic except RSA amplitude which returned to baseline value only 60 min following the stress test. We conclude that the neuronal network for OTergic amplification of RSA is used endogenously during recovery from stress, a calming behavior.

## Discussion

We have identified a multisynaptic neuronal network, from the hypothalamus to the brainstem to the heart, for OTergic amplification of RSA, and its implication in the recovery from stress (Extended Data Fig. 8). OT neurons in the caudal PVN in the hypothalamus project to the preBötC/nA and the DMV in the brainstem. PVN^OT^ fibers exert a dual control on HR *via* both areas, inducing RSA amplification through the release of OT in the preBötC/nA, and concomitantly decreasing mHR *via* the DMV. In the preBötC/nA, the OT-R is expressed in a subgroup of preBötC neurons. PreBötC^OT-R^ neurons are predominantly inhibitory (glycinergic and GABAergic), they regulate both respiratory and cardiac activities, forming a cardiorespiratory hub that controls RSA amplitude. PreBötC^OT-R^ neurons make putative monosynaptic contacts with nA^Cardiac^ neurons. OT increases the excitability of preBötC neurons, amplifying the inspiratory inhibitory (glycinergic) connectivity from preBötC to nA neurons. This induces an amplification of the respiratory modulation of cardiac parasympathetic activity, thereby amplifying RSA. On the other hand, sympathetic vasomotor activity is not affected. Our work has thus identified a PVN^OT^→preBötC^OT-R;Glycine^→nA^Cardiac^→Parasympathetic→Heart circuit for RSA amplification, which was found in female and male mice as well as in rats. This connectivity was identified on different experimental preparations in newborn, juvenile and adult animals. Behaviourally, OT does not influence RSA amplitude in resting adult animals, a condition where RSA amplitude is actually large. A stressful event, such as a restraint test, induces a large decrease in RSA amplitude, and OT neurons are involved in the rapid recovery in RSA amplitude during the recovery from stress, but not in the recovery of other respiratory and vascular parameters.

OT neurons were previously shown to regulate cardiac function, including by decreasing mHR, *via* an action on DMV^Cardiac^ neurons^23,26^. However, DMV^Cardiac^ neurons represent only 20% of all cardiac parasympathetic preganglionic neurons, and are commonly referred to as the vagal slow lane to the heart because their axons are unmyelinated C fibers in rodents. Our work showing that OT neurons can only evoke a small (< 5%) decrease in mHR is consistent with the bradycardia levels previously obtained upon direct stimulation of DMV neurons or regulation by OT. Conversely, nA^Cardiac^ neurons represent the remaining 80% of all cardiac parasympathetic preganglionic neurons, and are referred to as the vagal fast lane to the heart because their axons are myelinated B fibers. Stimulation of nA neurons induces fast and profound bradycardia, which is consistent with the encoding of RSA - i.e. fast and large oscillations in HR - *via* nA^Cardiac^ neurons. We show that OT can induce a large amplification of RSA (> 50%) by modulating the strength of the preBötC^OT-R;Glycine^→nA^Cardiac^ neuronal connectivity. Therefore, our data provide further evidence for the dichotomy in the regulation of cardiac activity by nA^Cardiac^ and DMV^Cardiac^ neurons, with OT neurons regulating independently RSA amplitude and mHR *via* each nucleus (respectively).

Our results also identify a phenotypic signature of key nodal neurons - preBötC^OT-R^ neurons - that control RSA amplitude by generating at least part of the inspiratory component of RSA. Optogenetic modulations of the activity of preBötC^OT-R^ neurons enabled large and bidirectional variations in RSA amplitude (+80 % / -50 %), indicating that this preBötC neuronal subgroup participates in setting the ongoing RSA level. Further, we show that (i) preBötC^OT-R^ neurons are predominantly inhibitory, (ii) they are presynaptic partners to nA^Cardiac^ neurons, and (iii) OT application enhances the excitability of preBötC inspiratory neurons and amplifies the inspiratory inhibitory (glycinergic) post-synaptic currents recorded in nA neurons. Given that about half of the inhibitory preBötC neurons are inspiratory^36,44^, collectively our data show that preBötC^OT-R^ neurons generate at least part of the inhibitory inspiratory modulation of cardiac parasympathetic activity, which leads to the increase in HR during inspiration. This is a major component in the generation of RSA, since HR automatically decreases after inspiration due to a predominant parasympathetic tone in mammals at thermoneutrality. PreBötC^OT-R^ neurons may even participate in the increased parasympathetic tone just after inspiration, by inducing a post-inhibitory rebound of excitation on nA neurons after inspiration (Fig. 4h-i). Still, part of the cardiac parasympathetic tone originates from respiratory neurons that are active during early expiration (post-inspiration), most likely the pontine Kölliker-Fuse neurons^42^. Future studies could investigate a putative role of OT in the amplification of the expiratory component of RSA.

When anxious, stressed or scared, and feeling our heart pounding in our chest, taking deep breaths is a universal way to calm down. Indeed, slow and deep breaths strongly amplify RSA and decrease mHR. Ancestral breathing techniques, such as yogic pranayama, and more recent ones like “boxed” breathing or the “365” breathing method, all rely on deep slow breathing to amplify RSA. Even Buddhist mantras and Catholic rosary prayers amplify RSA by inducing slow deep breathing^45^. In all these practices, voluntary deep slow breathing is used to indirectly decrease mHR, by amplifying the respiratory modulation of cardiac parasympathetic activity and therefore RSA. Our work identifies a non-voluntary hypothalamus-brainstem neuronal network that similarly amplifies RSA when calming down during recovery from stress. This network could participate more globally in the amplification of RSA during positive socio-emotional states in humans where breathing is not consciously altered, such as during parental bonding. Interestingly, the OTergic amplification of RSA is mediated by a specific amplification of the respiratory→cardiac parasympathetic (preBötC^OT-R;Glycine^→nA^Cardiac^) connectivity, with very little (if any) alteration in the respiratory activity *per se*. Therefore, RSA amplification by OT administration could be used as a therapeutic strategy in cardiovascular and mental disorders in which RSA amplitude is severely decreased, without adverse effects on respiratory function. Finally, whether variations in RSA amplitude are exclusively a physiological representation of mental states, or can also cause variations in mental states *per se*, remains unanswered. Recently, the finding that tachycardia is anxiogenic *per se*, which has been empirically accepted but never definitively proven, was demonstrated through elegant and selective optogenetic control of HR^46^. We propose that chronic modulations of preBötC^OT-R^ neurons will enable the selective amplification or reduction of RSA, to clearly resolve its physiological and psychological functions.

## Methods

**Table 1:** Animal sources.

| Species and genetic background | Designation | Reference | Additional informations |
| --- | --- | --- | --- |
| Mice, C57BL6/J | WT |  | Charles River |
| Mice, Swiss | WT |  | Janvier Labs |
| Mice, C57BL6/J | OT::cre | (Wu *et al.*, 2012)<sup>49</sup> | JAX stock #024234 |
| Mice, C57BL6/J | Ai27(LSL-ChR2) | (Madisen *et al.*, 2012)<sup>50</sup> | JAX stock #012567 |
| Mice, C57BL6/J | OXTR::cre | (Daigle *et al.*, 2018)<sup>51</sup> | JAX stock #031303 |
| Mice, C57BL6/J | Ai14(LSL-tdTomato) | (Madisen *et al.*, 2010)<sup>52</sup> | JAX Stock 007914 |
| Mice, mixed | GAD67::EGFP | (Tamamaki *et al.*, 2003)<sup>53</sup> | Gift from Dr. A.Baude, Inmed |
| Mice, C57BL6/J | GlyT2::EGFP | (Zeilhofer *et al.*, 2005)<sup>54</sup> | Zeilhofer lab, ETH Zurich |
| Mice, C57BL6/J | OT::GFP |  | GENSAT project: Tg(Oxt-eGFP)LC317Gsat/Mmucd |
| Mice, C57BL6/J | R26-LSL-hM4Di-DREADD | (Zhu *et al.*, 2016)<sup>64</sup> | JAX stock #026219 |
| Rat, Wistar | WT |  | Janvier Labs |

**Table 2:** Virus and tracer sources.

| Type | Designation | Source | Identifiers | Additional information |
| --- | --- | --- | --- | --- |
| Virus, AAV | AAV-phSyn1-FLEX-tdTomato-T2A-sypEGFP-WPRE | Addgene | Addgene #51509 | Titer : $1.8 \times 10^{13}$ GC/ml |
| Virus, AAV | AAV9-hSyn-DIO-somBIPOLES-mCerulean | Addgene | Addgene #3154951 | Titer : $1.7 \times 10^{13}$ GC/ml |
| Tracer | FluoroGold | Fluorochrome |  | 0.2 % (IP), 0.5 % (intracerebral) in NaCl |
| Tracer | Cholera toxin subunit B, Alexa Fluor™ 488 Conjugate (Ctb-488) | ThermoFisher | C34775 | 0.01% in NaCl |
| Tracer | Fluorescent beads (580/605 nm) | ThermoFisher | F8793 | 0.5 % in aCSF without D-Glucose |

**Table 3:**
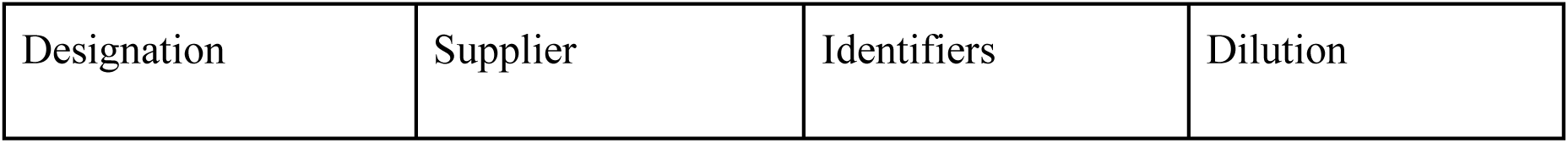
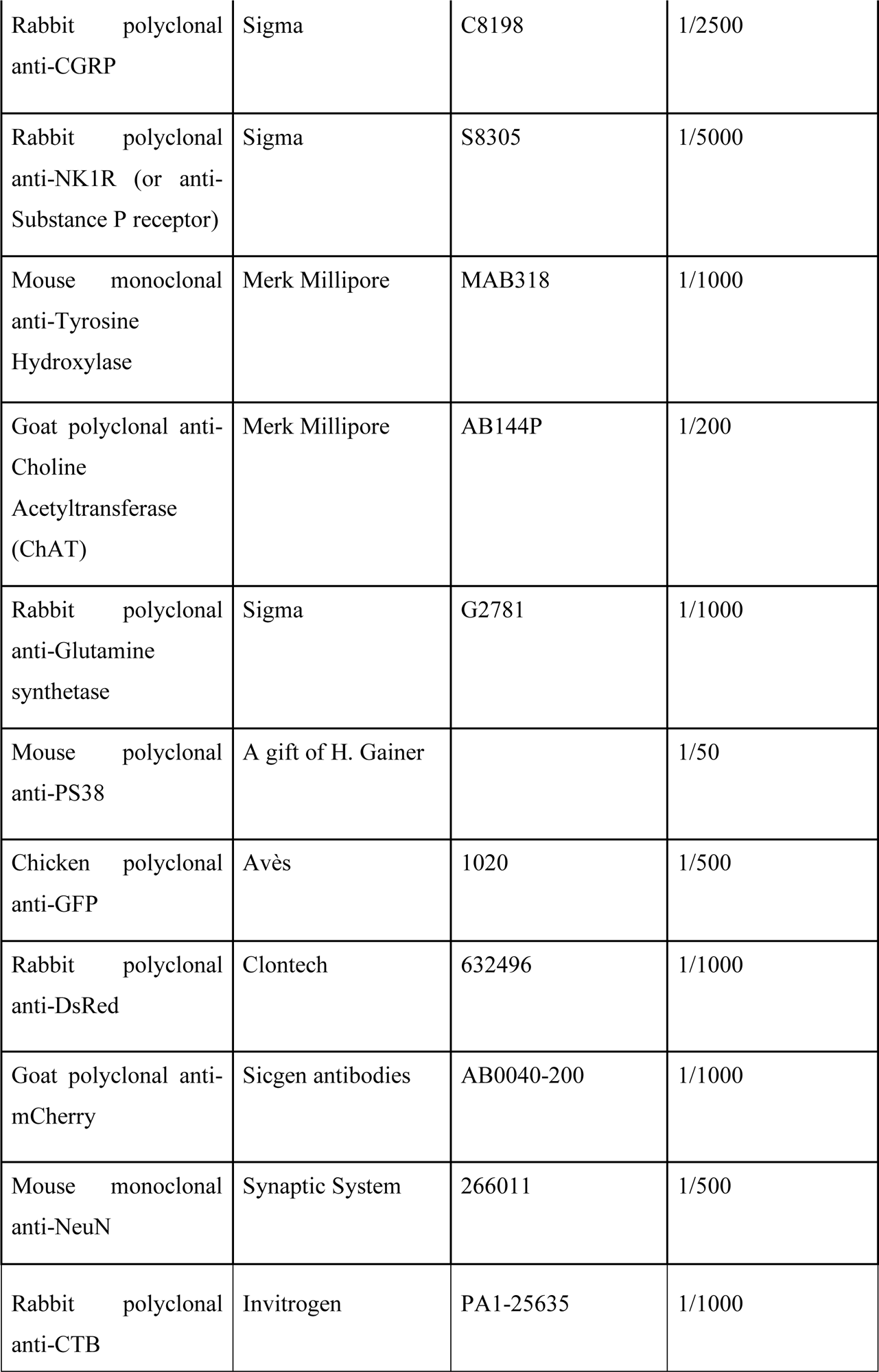
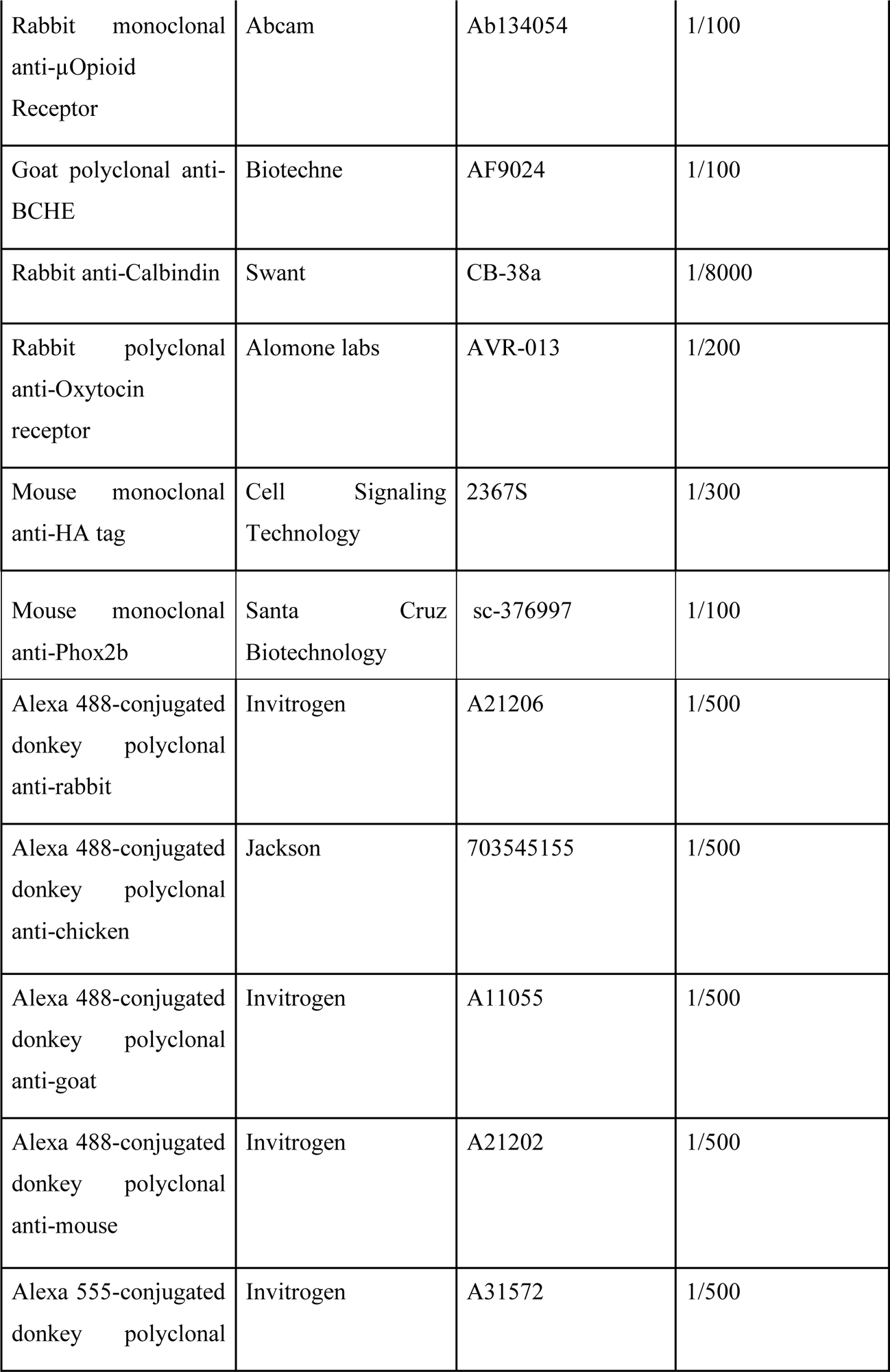
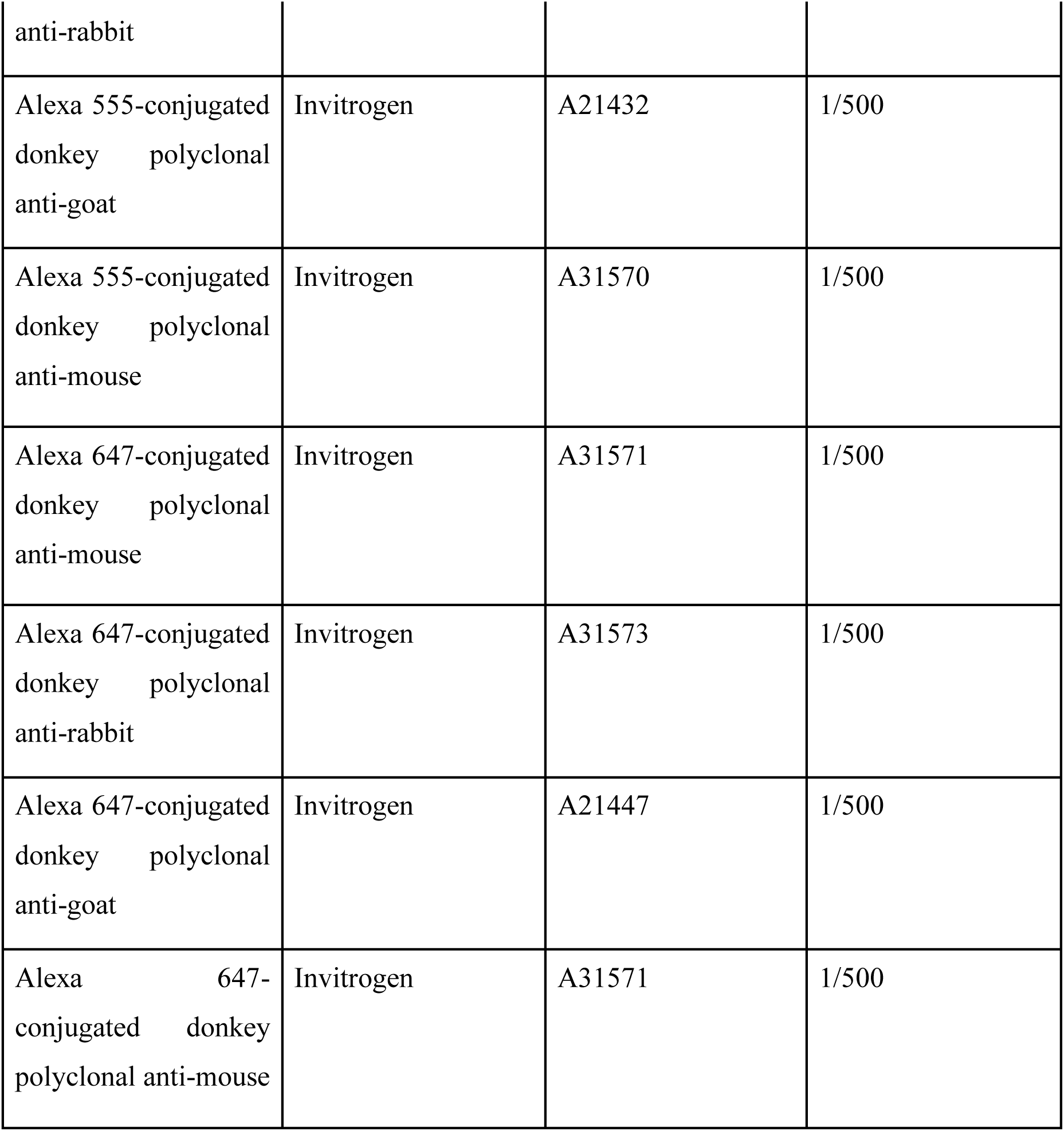
Primary and secondary antibody sources.

**Table 4:** Chemical compound sources.

| Designation | Supplier | Identifiers | Concentration |
| --- | --- | --- | --- |
| (Thr<sup>4</sup>,Gly<sup>7</sup>)-Oxytocin (TGOT) | Bachem | 4013837.0005 | 0.5 μM |
| (d(CH<sub>2</sub>)<sub>5</sub><sup>1</sup>,Tyr(Me)<sup>2</sup>,Thr<sup>4</sup>,Orn<sup>8</sup>,des<br>-Gly-NH<sub>2</sub><sup>9</sup>)-Vasotocin<br>trifluoroacetate salt | Bachem | 4031339 | 1 μM |
| Isoflurane | Axience | Isorane | 4 % induction, 2 % holding |
| Vasopressin acetate | Sigma | V9879 | 2 nM |
| Vecuronium bromide | Sigma | PHR1627 | 2 µg/ml |
| Cyanure (KCN) | Sigma | 1064371000 | 0.01 % |
| Atropine | Sigma | A0132 | 10 mg/kg (IP) |
| Ficoll 70 | Sigma | 17-0310-50 | 1.25 % |
| DREADD agonist 21 (Compound 21) | Hello Bio | HB6124 | 2.5 mg/kg (IP) |

### Animal experiments

Animals were handled and cared for in accordance with the Guide for the Care and Use of Laboratory Animals (N.R.C., 1996) and the European Communities Council Directive of September 22^th^ 2010 (2010/63/EU,74). Experimental protocols received approval from the institutional Ethical Committee guidelines for animal research from the French Ministry of Agriculture (authorization APAFIS #12006). Animals were group-housed with a 12-h light-dark cycle, at a constant temperature (22 ± 1°C) with *ad libitum* access to standard chow and water. Experiments were performed on transgenic or wild-type (WT) mice and rats (Table 1). All efforts were made to reduce animal suffering and minimize the number of experimental animals.

### Viruses

The AAV9-hSyn-DIO-somBIPOLES-mCerulean (somBiPOLES) virus for optogenetic inhibition or excitation of preBötC^OT-R^ neurons in OT-R::Cre mice was obtained from Addgene (Table 2, Addgene viral prep # 154951; http://n2t.net/addgene:154951; RRID:Addgene_154951)^32^. The AAV1-phSyn1(S)-FLEX-tdTomato-T2A-SypEGFP-WPRE virus for tracing experiments from preBötC^OT-R^ neurons to nA^Cardiac^ neurons in OT-R::Cre mice was obtained from Addgene (Table 2, Addgene viral prep # 51509; http://n2t.net/addgene:51509; RRID:Addgene_51509)^55^.

### Viral and tracer microinjections in the preBötC

For tracing experiments, 3/4-month-old male and female mice received a pre-surgery injection of a non-steroidal, anti-inflammatory drug (carprofen, Rimadyl^®^, 5 mg/kg, s.c.). Anesthesia was induced by inhalation of isoflurane (4 %, 80 % O_2,_ Isorane, 1000 mg/g, Axience, France) in an induction box. Mice were placed in a stereotaxic frame with the nose ventro-flexed (40°) and maintained under isoflurane anesthesia (2 %). Body temperature was maintained at 37.5°C using a heating pad (40-90-8D model, FHC, ME, USA). A midline incision of the skin and muscle tissues was made at the occipital level, a cranial window was created on the caudal occipital bone using a dental drill, and the atlanto-occipital membrane was opened to expose the brainstem. Glass micropipettes (∼30 µm tip diameter) were filled with injectate and connected *via* a silver wire to an electrophysiology amplifier (Neuro Amp Ex, ADInstruments) for neuronal activity recording. The pipette (20° angled in the rostro-caudal axis, tip pointing forward) was descended into the brainstem 1.25 mm lateral to the midline. Extracellular recordings of multiunit activity were used to functionally map the preBötC, defined as the most rostral point displaying distinct inspiratory-locked activity (typically 0.0 ± 0.2 mm rostral, 1.4 ± 0.2 mm ventral to the *calamus scriptorius*).

For the somBiPOLES virus injections, the mice were placed in the stereotaxic frame with a horizontal head position. The extracellular recording pipette was tilted 10° in the rostro-caudal axis, pointing towards the caudal part of the mouse. A cranial window was made at the rostral end of the occipital bone, at its junction with the parietal bone. Glass micropipettes were descended through the cerebellum to the brainstem, 1.25 mm lateral to the midline. Injection spots were typically 1.7 ± 0.3 mm caudal and 6.0 ± 0.3 mm ventral to the *Lambda*.

Bilateral injections were performed for all experimental conditions, except for unilateral FluoroGold injections. Viral vectors (25 nl for the tracing virus and 35 nl for the somBiPOLES virus per injection site over 5 min, into two rostro-caudal sites 200 µm apart) or tracers (Cholera toxin subunit B (CTB): 70 nl per injection site over 5 min, into three rostro-caudal sites 200 µm apart; FluoroGold: 25 nl per injection site over 5 min, into two rostro-caudal sites 200 µm apart) were microinjected using a picospritzer (PICOSPRITZER^®^ III, Parker, NH, USA) and a graduated monocular. At each site, pipettes were left in place for 10 min after injection before being withdrawn. Wounds were closed with sterile sutures. Experiments were conducted 3 weeks after viral injections and 1 week after tracer injections. All injection sites were verified following the experiments.

### Optogenetic cannula and ECG telemetric transmitter implantation

For *in vivo* freely moving experiments, OT::cre;Ai27(LSL-ChR2) mice (3/4-month-old) were anesthetized and placed in a stereotaxic frame as described above for preBötC microinjections. A midline incision was made over the cranial bone and all attached muscles were removed. Head horizontality was checked using *bregma* and *lambda* coordinates, and a cranial window was made at the rostral level of the occipital bone, at the junction with the parietal bone. ECG and EMG inspiratory recordings were obtained using 3 electrodes: two in cranial muscles and one subcutaneously through a dorsal incision, which was subsequently used to insert the telemetric transmitter. Signals were amplified, filtered, digitized (Power 1401, Cambridge Electrical Design, Cambridge, UK), rectified and integrated by Spike2 software (Cambridge Electrical Design, Cambridge, UK). Extracellular recordings were performed to map the preBötC, following the same techniques as for somBiPOLES viral injections (typically 1.25 mm lateral to the midline and 1.2 ± 0.3 mm caudal, 6.0 ± 0.3 mm ventral to the *Lambda*). Bilateral optogenetic cannulas (200 µm, NA 0.22, Doric Lenses Inc., Québec, Canada) were then implanted at the preBötC rostro-caudal and medio-lateral coordinates. The dorso-ventral position was determined based on optimal RSA amplification and bradycardia induction during optogenetic stimulation, typically 0.4 mm above the best extracellular inspiratory recording. Cannulas were fixed with cyanoacrylate (Super Glue, Loctite).

During the same surgery, ECG telemetric transmitters (ETA-F10, Data Sciences International (DSI), Harvard Bioscience, Inc., MN, USA) were also implanted. The transmitter body was placed subcutaneously along the mouse’s flank, using the dorsal incision made at the beginning of the surgery. Two subcutaneous electrodes, previously exposed by 1 cm, were fixed with sutures near the right shoulder and the left rib region. The wound was closed with sterile sutures. ECG signal from the transmitter was recorded (Matrix 2.0, DSI, Harvard Bioscience, Inc., MN, USA), sampled at 2000 Hz, and integrated into Spike2 software using a DSI talker.

After a 1-week recovery, respiratory activity was recorded by plethysmography and ECG by telemetry for 3h, during which optogenetic stimulations of OT^+^ fibers in the preBötC were performed periodically.

### *In vivo* plethysmography recordings

Breathing of unrestrained, non-anesthetized mice was recorded using whole-body plethysmography (EMKA Technologies, Paris, France). A constant air flow (0.5 l/min) circulated inside the 200 ml plethysmography chambers (maintained at 25 ± 0.5 °C) using a vacuum pump (Vent 4, EMKA, Paris, France). During each recording, the amplitude of the signal measured was calibrated by injecting 1 ml of air into the chamber containing the mouse. Analog signals were amplified (Amplipower, EMKA, Paris, France), converted (Power 1401, Cambridge Electrical Design, Cambridge, UK) and integrated into Spike2 software (Cambridge Electrical Design, Cambridge, UK).

### *In vivo* RSA recordings in anesthetized animals

For RSA recordings in the anesthetized condition *in vivo*, animals were anesthetized and placed in a stereotaxic frame similar to preBötC microinjections for tracing experiments, with the nose ventro-flexed (40°), providing access to the brainstem surface. For mice experiments, ECG and EMG inspiratory recordings were obtained using 3 electrodes placed under the skin of the hind legs and a front leg. For rat experiments, the EMG inspiratory recordings were obtained *via* external intercostal muscle electrodes and the ECG was recorded using three electrodes placed under the skin of the hind legs and thorax. All signals were then amplified, filtered, digitized (Power 1401, Cambridge Electrical Design, Cambridge, UK), rectified and integrated by the Spike2 software (Cambridge Electrical Design, Cambridge, UK).

In Wistar rats, as described previously, extracellular recordings were made to functionally map the preBötC bilaterally (typically 1.8 mm lateral to the midline and 0.7 ± 0.2 mm rostral, 1.7 ± 0.3 mm ventral to the *calamus scriptorius*). A specific OT-R agonist (TGOT^37^, 0.5 µM diluted in artificial cerebrospinal fluid (aCSF, composition as used for the Working Heart-Brainstem preparation below), 50 nl) or aCSF (vehicle injections, 50 nl), diluted with 0.5% fluorescent beads, was slowly injected bilaterally into each preBötC using a syringe and a graduated monocular.

In OT::cre;Ai27(LSL-ChR2) mice, unilateral photostimulations were performed in the preBötC. The optical fiber was tilted by 20° in the rostro-caudal axis, with the tip pointing towards the animal’s head. The preBötC was located by the following coordinates: 1.25 mm lateral to the midline and 0.0 mm rostral to the *calamus scriptorius*. The dorso-ventral coordinate was functionally determined based on the optogenetic stimulation-induced bradycardia and RSA amplification. After identifying the optimal position, several stimulations were performed at intervals of at least 5 min. To precisely perform OT-R antagonist injections into the preBötC, extracellular recordings were made, as previously described. Optimal inspiratory extracellular activities were generally located 400 µm ventral to the coordinate defined for the optogenetic stimulations. OT-R antagonist (d(CH_2_)_5_^1^,Tyr(Me)^2^,Thr^4^,Orn^8^,des-Gly-NH_2_^9^)-Vasotocin trifluoroacetate salt (OTA), 1 µM diluted in aCSF, 200 nl, Table 4) was then injected unilaterally in the preBötC^23,25,56^. Several photostimulations were performed every 5 min for 1h after OTA injection. In some animals, photostimulations were also performed on the contralateral preBötC and in the ipsilateral DMV (coordinates: 0.3 mm lateral to midline, 0.0 ± 0.3 mm rostral and 0.2 ± 0.2 mm ventral to the *calamus scriptorius*). For control experiments aCSF without OTA was injected. Fluorescent beads (0.5%) were added to all injectate solutions to enable the localisation of the injection spots *a posteriori*.

In mice injected with the somBiPOLES virus, bilateral photoexcitations and photoinhibitions were performed in the preBötC using the techniques described for photostimulations in OT::cre;Ai27(LSL-ChR2) mice experiments. At the end of the experiment, intraperitoneal injection of atropine was administered (10 mg/kg, diluted in NaCl, 600 µl for a 30g mouse, Table 4), followed by additional stimulations after a 20-min interval.

### Optogenetic stimulations

For the experimental setups, uni- or bi-lateral cannulas (200 µm, NA 0.22, Doric Lenses Inc., Québec, Canada) were connected to one or two optical fibers (200 µm, NA 0.22, Doric Lenses Inc., Québec, Canada), themselves connected to a rotary joint allowing the laser light beam to be split if necessary. A common optical fiber was connected to either a blue laser (473 nm, DPSS, ADR-800A, Shanghai Laser & Optics Century Co., Ltd., China), or a dual laser with blue and red light (473 nm and 561 nm, DPSS, 06 Series, Cobolt AB, Hübner Photonics GmbH, Germany) for mice injected with the somBiPOLES virus. Lasers were triggered by a TTL-Pulse-generator (PulserPlus, Prizmatics Ltd., Israel) configured in Pulser software and signals were integrated into Spike2 software.

In experiments involving OT::cre;Ai27(LSL-ChR2) mice, either in freely moving or anesthetized conditions, photostimulations were conducted using 10 ms pulses at 10mW/mm² power, 3000 pulses, and a frequency of 50 Hz. Photostimulations in freely moving experiments were performed during calm periods characterized by a stable respiratory frequency.

In experiments with mice injected with the somBiPOLES virus, photoinhibition was carried out using the blue laser (473 nm) and consisted of 10 ms pulses (1.2 mW/mm², 3000 pulses, 50 Hz), whereas photoexcitation was performed with the red laser (561 nm) and consisted of 10 ms pulses (10 mW/mm², 3000 pulses, 50 Hz). The power difference was due to the varying sensitivities of the two opsins.

A minimum of 5 min was allowed between two consecutive photostimulations in all experiments. A light meter (PM100D fiber power meter, Thorlabs, Newton, NJ, USA) was used to calibrate the light density imposed by each optical fiber up to 10 mW/mm² (for a distance of around 400 µm between the optical fiber and the preBötC/nA), as determined to induce maximal effects without any adverse impact on the tissue^57^. All photostimulation protocols were repeated multiple times in each animal and a strong intra-preparation reproducibility was noted.

### Working Heart-Brainstem preparation

Experiments were performed using the arterially perfused *in situ* Working Heart-Brainstem preparation (WHBP)^13,58^, as previously described^12,40^, and used on juvenile Wistar rats of both sexes, aged 21-30 days. Rats were deeply anesthetized with isoflurane, bisected below the diaphragm, exsanguinated and cooled in aCSF solution on ice (composition in mM: 125 NaCl, 24 NaHCO3, 3 KCl, 2.5 CaCl2, 1.25 MgSO4, 1.25 KH2PO4 and 10 D-Glucose, pH 7.3 after saturation with carbogen gas (chemicals were purchased from Sigma-Aldrich, France)). The decerebration was performed precollicularly, the lungs were removed, and the descending aorta was isolated and cleaned. Preparations were transferred to a recording chamber. Retrograde perfusion of the thorax and head was achieved via a double-lumen catheter (ø1.27 mm, PowerPICC, Bard, NJ, USA) inserted into the descending aorta. The perfusion solution consisted of aCSF solution containing Ficoll (1.25 %), vasopressin acetate (2 nM, V9879, Sigma-Aldrich, France) and vecuronium bromide (2-4 µg/ml, PHR1627, Sigma-Aldrich, France) warmed to 31°C and gassed with carbogen (closed loop reperfusion circuit, 25-30 ml/min). A transducer connected to the second lumen of the catheter monitored perfusion pressure (PP) in the aorta. The head of the preparation was fixed with ear and mouth bars. A peristaltic pump (520S, Watson-Marlow Bredel Pumps, UK) was used to slowly increase the flow rate of the perfusion solution to tune the preparation. For most WHBPs, simultaneous recordings of phrenic nerve activity (PNA), cervical vagus nerve activity (VNA, left side) and thoracic sympathetic chain activity (tSNA, between T8-10) were obtained using glass suction electrodes.

For some WHBPs the cardiac vagal branch activity (CVBA) was recorded, instead of the cervical vagus nerve activity, in accordance with the protocols described previously^42^. Briefly, a small amount of parenchymal tissue around the cut left bronchus was kept and the thymus was resected to gain access to the left cardiac vagal branch. In the recording chamber, the left thoracic vagus nerve could be observed parallel to the oesophagus and was carefully dissected away from the surrounding tissue to expose its branches. The left cardiac vagus branch was usually found just caudal to the left recurrent laryngeal nerve and was cut as distally as possible. The functional validity of the cardiac vagal branch recording was tested with an excitatory baroreceptor challenge by briefly (2-3 s) increasing the perfusion pump setting to its maximum, taking care that rising fluid levels caused no artefact, and chemoreceptor challenge by injection of KCN (0.2 ml, 0.01 %). Preparations that failed this test were either repositioned until the test was successful or discarded.

All neurograms were amplified, filtered (300-5000 Hz band-pass, A-M system, 1700 Model, Carlsborg, USA), digitized (Power 1401, Cambridge Electrical Design, Cambridge, UK), rectified and integrated using Spike2 software (Cambridge Electrical Design, Cambridge, UK). Heart Rate (HR) was derived by using a window discriminator to trigger from the systolic pressure recorded from the perfusion pressure sensor.

Once the WHBP was tuned, bilateral brainstem extracellular recordings were performed to locate inspiratory activity from preBötC neurons. They were typically found 1.5mm lateral, 0.9 ± 0.3mm rostral, 2.0 ± 0.3 mm ventral to the *calamus scriptorius*. TGOT (0.5 µM, 50 nl, diluted in aCSF with 0.5% fluorescent beads) or aCSF (vehicle injections, 50 nl, with 0.5% fluorescent beads) were slowly injected in each preBötC bilaterally using a syringe and a graduated monocular.

Nerve activity and HR traces were smoothed before analysis (PNA: time constant 0.05 s (DC remove); tSNA: time constant 0.1 s (smoothing) and 0.032 s (DC remove); VNA: time constant 0.2 s (smoothing) and 0.032 s (DC remove); CVBA: time constant 0.032 s (DC remove); HR: time constant 0.5 s (smoothing)).

### Rhythmic preBötC/nA slice and island preparations

Brainstem transverse slice preparations containing the preBötC network and the nA were obtained from mouse pups (P0-P5) as follows. Swiss newborns were quickly decapitated at the C4 spinal level and the brainstem was carefully dissected in a chilled oxygenated aCSF composed of (in mM): 120 NaCl, 8 KCl, 1.26 CaCl2, 1.5 MgCl2, 21 NaHCO3, 0.58 NaH2PO4, 30 glucose, pH 7.4. The isolated hindbrain was embedded in a low melting point agar block (ThermoFisher, France) and oriented to enable serial transverse sectioning along the rostro-caudal axis using a vibratome (VT1200S, Leica, Germany). A 500 µm thick slice, with its anterior limit set ∼300 µm caudal to the posterior limit of the facial nucleus, was isolated. To define the correct rostro-caudal level at which the expected slice would be obtained we also used other anatomical landmarks as previously described^59^. Slices were then transferred to a recording chamber, rostral surface upwards, and continuously superfused with oxygenated aCSF at 30°C. Preparations were allowed to recover for 20-30 min before any recording sessions began.

The island preparations were obtained from transverse slices obtained as detailed above by additionally trimming away tissue outside the region hosting the preBötC network and the nA using fine scissors^60^.

### Procedure for recordings in preBötC/nA slice and island preparations

PreBötC network activities in slice and island preparations were recorded using glass micropipettes (tip diameter 80-100 µm) positioned on the slice’s rostral surface, ventral to the nA where the preBötC respiratory circuitry is located. The micropipettes were fabricated from aCSF-filled borosilicate glass tubes (Harvard Apparatus, Germany) broken at the tip and used as suction electrodes connected to a high-gain amplifier (AM Systems, USA). Signals were filtered (3Hz-3kHz), integrated (time constant 100 ms; Neurolog, Digitimer, England) and stored on a computer through a Digidata 1440 interface and PClamp10 software (Molecular Devices, USA).

Whole cell patch-clamp recordings were visually guided using differential interference contrast. For nA neurons recordings, the patch pipette was positioned close-to the nA pars compacta, slightly dorso-lateral to the micropipette monitoring preBötC activity. The patch pipette for recording individual preBötC neurons was also positioned close to the pipette used for extracellular recordings, and the recorded neurons were selected according to their activity in phase with the preBötC network activity. Patch pipettes were fabricated with borosilicate glass tubes using a puller (Sutter Instrument, USA) and filled with a solution composed of (in mM): 140 K-gluconate acid, 1 CaCl2.6H2O, 10 EGTA, 2 MgCl2, 4 Na2ATP, 10 HEPES, pH 7.2, supplemented with AlexaFluor 568 and had tip resistances of 5-7MΩ when filled with this solution. Electrophysiological signals were recorded using an Axoclamp 2A amplifier (Molecular Devices) and the same digitizing interface and software mentioned before. nA neurons were selected for patch-clamp recording on the basis of their location (between the dorsal part of the preBötC and the ventral part of the nA pars compacta) and their rhythmic inhibitory synaptic inputs received during each preBötC respiratory burst. The magnitude of the synaptic drive, representing the summation of inhibitory post-synaptic currents during individual inspiratory bursts, was measured as the maximal envelope amplitude, averaged over 10 consecutive bursts per neuron in slices.

Drugs were applied in the bath at a final concentration of: 0.5 µM TGOT (Bachem, Bubendorf, Switzerland); 10 µM Bicuculline (Trocris, Bio-Techne, MN, USA); 5 µM Strychnine (SigmaAldrich, St Louis, USA); 20 µM CNQX (SigmaAldrich, St Louis, USA).

### PVN slice preparations

P20 OT::Cre;Ai14(LSL-tdTomato);R26-LSL-hM4Di-DREADD mice were anesthetized using 5% isoflurane and rapidly decapitated. Coronal slices containing the PVN (350 µm) were obtained using a vibratome (VT 1200S, Leica, Germany). Slices were allowed to equilibrate before use for 1h in standard recording aCSF (in mM: 124 NaCl, 3 KCl, 1.2 NaH2PO4, 1.2 MgSO4, 25 NaHCO3, 11 D-glucose, and 2 CaCl2, saturated with 95% O2/5% CO2, pH 7.4, ∼300 mosM) at room temperature. During experiments the recording chamber was superfused at ∼3 mL/min with standard recording aCSF.

### Procedure for recordings in PVN slices

Electrodes (resistance ∼3.0-6.0 MOhm) were prepared using a Narishige micropipette puller (PC-100) and filled with (in mM) 130 K^+^-gluconate, 10 NaCl, 11 EGTA, 1 CaCl2, 10 HEPES, 1 MgCl2, 2 Mg-ATP, 0.2 Na-GTP, pH 7.3, and ∼280 mosM. Recording pipettes were guided using a piezoelectric manipulator (Luigs & Neumann, GmbH). PVN^OT^ neurons expressing tdTomato were identified in Cre^+^ mice using a Hamamatsu scientific camera. In Cre^-^ mice, unlabeled neurons in caudal PVN slices were recorded. Following initial membrane rupture the resting membrane potential was measured. Resting membrane potential, spontaneous action potentials, and evoked action potentials were recorded in current clamp mode. The DREADD agonist C21 (10 µM) was applied for 5 min. Data obtained from PVN slice experiments were acquired in pClamp 10.7 using a HEKA EPC 10 Double amplifier (HEKA Elektronik), filtered at 2 kHz and sampled at 20 kHz. Data were analyzed using Clampfit 10.7 and MiniAnalysis (Synaptosoft, USA).

### Histology and image acquisition

After *in vivo* anesthetized and WHBPs experiments, brains were fixed in antigenfix solution (Diapath, P0016, Italy) at 4°C for 48-72h and cryoprotected in 30 % sucrose. After *in vitro* slice experiments, the slices were fixed in 4 % paraformaldehyde for 2-3h and cryoprotected in 20 % sucrose solution. For other conditions, animals were transcardially perfused with 20 ml phosphate-buffered saline followed by 25 ml of antigenfix. Brains were removed, post-fixed in antigenfix at 4°C for 24h and cryoprotected in 30 % sucrose. In experiments requiring nA labeling, animals received an intraperitoneal injection of the retrograde tracer FluoroGold (0.2 %, 300 µl, Fluorochrome LLC, CO, USA) 5 days prior to perfusion. Coronal sections of brainstems or hypothalamus were obtained using a cryostat (CM350S, Leica Microsystems), with 30 µm sections for *in vitro* slices and 50 µm sections for all others.

For immunohistochemistry, 50 µm sections were blocked and permeabilized in phosphate-buffered saline (PBS, 1X) containing 5% normal goat serum, 2 % BSA and 0.3 % Triton X-100 for 1h at room temperature. 30 µm sections were incubated in PBS solution with 1 % BSA and 0.3 % Triton X-100 for 90 min. Sections were incubated with primary antibodies for 72h at 4°C (50µm sections) or overnight at room temperature (30 µm sections). After washing, sections were left for 2h (50 µm sections) or 90 min (30 µm sections) with secondary antibodies at room temperature. Sections were mounted on microscope slides using Vectashield mount medium without DAPI (H-1000, VectorLabs Inc., CA, USA) for FluoroGold-containing sections or with DAPI (H-1200, VectorLabs Inc., CA, USA) for others. Primary and secondary antibodies used are listed in Table 3.

Fluorescent staining was examined under a Zeiss LSM-800 (or LSM-900) confocal microscope with ×10, ×20 or ×40 objectives and captured using ZEN 2.6 (blue edition) imaging software. For cell counting analysis, z stacks images with 0.5 µm z-steps were used.

### Mapping of injection spots

For all experiments with chemical compound injections in the preBötC, fluorescent beads (excitation 580 nm, emission 605 nm, F8793, ThermoFisher Scientific Inc., MA, USA) were co-injected to enable the localisation of the injection spots. In each animal, sections were observed under a stereo-microscope (Olympus SZX16 with fluorescent lamp X-cite Series 120) and sections containing fluorescent beads were co-labeled with TH, NK1R antibody to localize preBötC, as well as ChAT antibody or Fluorogold (0.2 %, IP) to visualize nA. These markers helped align the sections in the Paxinos-Watson atlas (rats) or in the Paxinos-Franklin atlas (mice). Injection spots were imaged using an Apotome Zeiss epi-fluorescence microscope with a ×10 air objective and captured with an axioCam MRm camera and ZEN 2.6 imaging software (blue edition).

### Tissue clearing

Paraformaldehyde-fixed whole OT::GFP mouse brains were cut with a slicer to obtain coronal slices of ∼1mm thickness. Tissue was cleared using the CUBIC method^61–63^. Briefly, fixed tissue was immersed in 50%-diluted reagent-1A (containing 5 % aminoalcohol #10, 10 % Triton and 10 % urea) for 6h at 25°C with shaking, then replaced by reagent-1A for 42h at 25°C with shaking. From day2 to day4 tissue was shaken at 37°C. Clearing was stopped with PBS. The sliced tissue was then immersed with 50%-diluted reagent-2 (containing 50 % sucrose, 25 % urea, 10 % aminoalcohol #16 and 0.1 % Triton) for 24h at 25°C and then in reagent-2 for 48h at 25°C. The sliced tissue was mounted with a coverslip immersed in a 1:1 mixture of silicon oil and mineral oil. Images were acquired using a Zeiss LSM-800 confocal microscope.

### Blood pressure telemetric transmitter implantation

Male OT::cre;R26-LSL-hM4Di-DREADD mice were implanted with a telemetric blood pressure sensor (PA-C10, DSI, Harvard Bioscience, Inc., MN, USA) in the left carotid artery. 3/4-month-old mice were administered a non-steroidal anti-inflammatory drug (carprofen, Rimadyl^®^, 5 mg/kg, s.c.) 30 min prior to surgery. Anesthesia was induced with isoflurane inhalation in an induction box, followed by an intraperitoneal injection of a ketamine-xylazine mixture (ketamine: 100 mg/kg, Imalgene 1000, IP; xylazine: 10 mg/kg, Rompun 2%, IP). Mice were positioned on their back on a heating pad maintaining their temperature at 37.5°C. A midline neck incision was made, and the carotid artery sheath was accessed by gently displacing the salivary glands and muscles. The left carotid artery was isolated from the vagus nerve and carotid vein until the carotid bifurcation was located. A ligature was placed around the carotid bifurcation and used to create tension, facilitating the positioning of a vascular clamp approximately 1 cm upstream of the bifurcation. A needle was used to make an incision in the carotid artery, allowing catheterization with the telemetric sensor catheter. Once in place, the clamp was removed, and the catheter was inserted approximately 1 mm into the aorta and ligated to the carotid artery. The transmitter was subcutaneously inserted into the back of the animal. The incision was closed using sterile sutures, and lidocaine (5% cream, Lidocaïne Prilocaïne, Biogaran, France) was applied to the incision area. The animal was allowed to recover for 1 week.

### Behavioral studies of recovery from stress

Prior to chemogenetic experiments, male OT::cre;R26-LSL-hM4Di-DREADD mice were habituated daily to their plethysmography chamber for 1 week. Then, mice received a slight isoflurane anesthesia for intraperitoneal injection of compound 21 (2.5 mg/kg diluted in NaCl 0.9 %, DREADD agonist 21 dihydrochloride, HB6124, Hello Bio Ltd, Ireland) or vehicle solution (NaCl 0.9 %). The mice were then placed in their plethysmography chamber, which was positioned on the receiver plate to simultaneously record respiratory activity and blood pressure. Respiratory signals were amplified (Amplipower, EMKA, Paris, France) and digitized (Power 1401, Cambridge Electrical Design, Cambridge, UK). Blood pressure telemetry signals were recorded (Matrix 2.0, DSI, Harvard Bioscience, Inc., MN, USA) and rectified according to the measurement of ambient pressure (APR-2, DSI, Harvard Bioscience, Inc., MN, USA). Respiratory and telemetric signals were integrated into Spike2 software (Cambridge Electrical Design, Cambridge, UK) at 2000 Hz sampling rate. After a 1-hour control period in a low-light room at room temperature, mice were placed in a restraining ventilated cylinder that severely restricted their movements for 10 min, before returning to the plethysmography chamber for 70 min. Heart rate was derived from systolic blood pressure through automatic detection threshold with manual verification. Analysis was performed before the stress stimulus (during calm periods), during the stress phase, and twice during the post-stress recovery phase (25-35 min and 55-65 min post-stress). Stress effects on respiratory parameters and RSA were quantified within the first 5 min after mice returned to their plethysmography chamber. Each animal had six recording sessions over consecutive days, lasting about 2h30 each, to obtain three recordings in each condition (Compound 21 or NaCl). NaCl sessions always preceded Compound 21 sessions due to the compound’s prolonged effects (up to 6 hours).

### Data analysis and statistics

Cell counting was carried out on serial 1:4 sections, following the imaging process previously described. Cell counting of OT-R^+^, Calbindin^+^ and BCHE^+^ neurons in the preBötC/nA, as well as quantifications in OT-R::Cre;Ai14(LSL-tdTomato);GAD67::GFP mice and OT-R::Cre;Ai14(LSL-tdTomato);GlyT2::GFP mice, were performed on serial 1:2 sections. Manual counting was performed unilaterally. For PVN counts, three sections were selected to cover the entire rostro-caudal axis. For nA counts, all sections with FluoroGold injection or ChAT labeling were included, and all labeled sections were counted (approximately 5-6 sections per series). PreBötC counts were based on the location of the nA, ventricular aperture, slice shape, and marker selection, resulting in a set of slices containing preBötC neurons (approximately 3 slices per series). The total number of each cell type per group was estimated by the sum of the neurons counted per series, multiplied by 4 (series 1:4) or by 2 (series 1:2).

Measurements of RSA amplitude were always performed following respiratory triggered averaging of HR. The trigger was placed at the end of each inspiration, for >15 respiratory cycles in each condition studied. HR windows corresponding to each respiratory cycle centered on the end of inspiration were then averaged to enable an accurate measurement of mean RSA amplitude in each condition studied.

Delta values in percentages (%) show the change in post-stimulus values relative to pre-stimulus values. Statistics on delta values to determine the effect of a given stimulus were performed on absolute values (raw data), when comparing the pre-stimulus and stimulus conditions. Group data are presented as mean ± SEM when in numerical values, and as violin plots (truncated or untruncated) with median and quartiles on graphs. Normal distribution of the data was tested with the Shapiro-Wilk test, and statistical analyses were performed using parametric or non-parametric tests accordingly. The statistical tests used and post-hoc analysis performed are detailed in the corresponding figure legends. All statistical analyses were performed using GraphPad Prism (v. 9.5.1, GraphPad Software, San Diego, CA). Statistical significance was set at P < 0.05. P values are indicated in brackets when between 0.05 and 0.1, but not for values above 0.1.

## Acknowledgements

We thank Dr Jean-Charles Viemari for assistance with preliminary rhythmic slice recordings, Laura Cardoit for assistance in immunostaining and confocal imaging of brainstem transverse slices, Dr Gilles Fortin for providing experimental equipment, Pr Andrew Allen and Dr John Simmers for their critical analysis of the manuscript, Dr Agnès Baude for providing the GAD67::GFP mouse line, the CTB tracer and the NeuN antibody, and members of the Inmagic, PBMC and Animal core facilities at Inmed. This work was supported by grants from the Agence Nationale de la Recherche (ANR-21-CE14-0009-01 to C.M.), from the French government under the Programme « Investissements d’Avenir » (Initiative d’Excellence d’Aix-Marseille Université via A*Midex funding (AMX-19-IET-004), and ANR (ANR-17-EURE-0029), to C.M.), from the French Ministry of Research (PhD fellowship to J.B.), from the Fondation pour la Recherche Médicale (FRM end-of-PhD Fellowship to J.B.), from Marseille Institute for Rare Diseases (Marmara, Master’s Fellowship to A.L.), and from the Institute Marseille Imaging (A*Midex AMX-19-IET-002).

## Author contributions

C.M. and J.B. designed the research project, with help from F.M. on matters related to oxytocin, and help from M.T.B. on in vitro electrophysiological aspects. J.B. performed all experiments except those cited hereinafter. C.M. performed the virus and tracer injections, and the optogenetic cannula implantations. M.T.B. performed the in vitro electrophysiology experiments on preBötC/nA rhythmic slices and their analysis. A.L. performed the in vivo anesthetized rat experiments and their analysis. R.T. performed the in vitro electrophysiology experiments on OT neurons and their analysis. C.G. and F.S. contributed to the histological experiments. M.S.F. and V.M. performed the CUBIC-clearing experiments. J.B. and C.M. analyzed the data. C.M. and J.B. wrote the paper.

## Data availability

The data are available upon request to the corresponding author. Source data are provided with this paper.

## Extended data figures and legends

**Extended Data Fig. 1:**
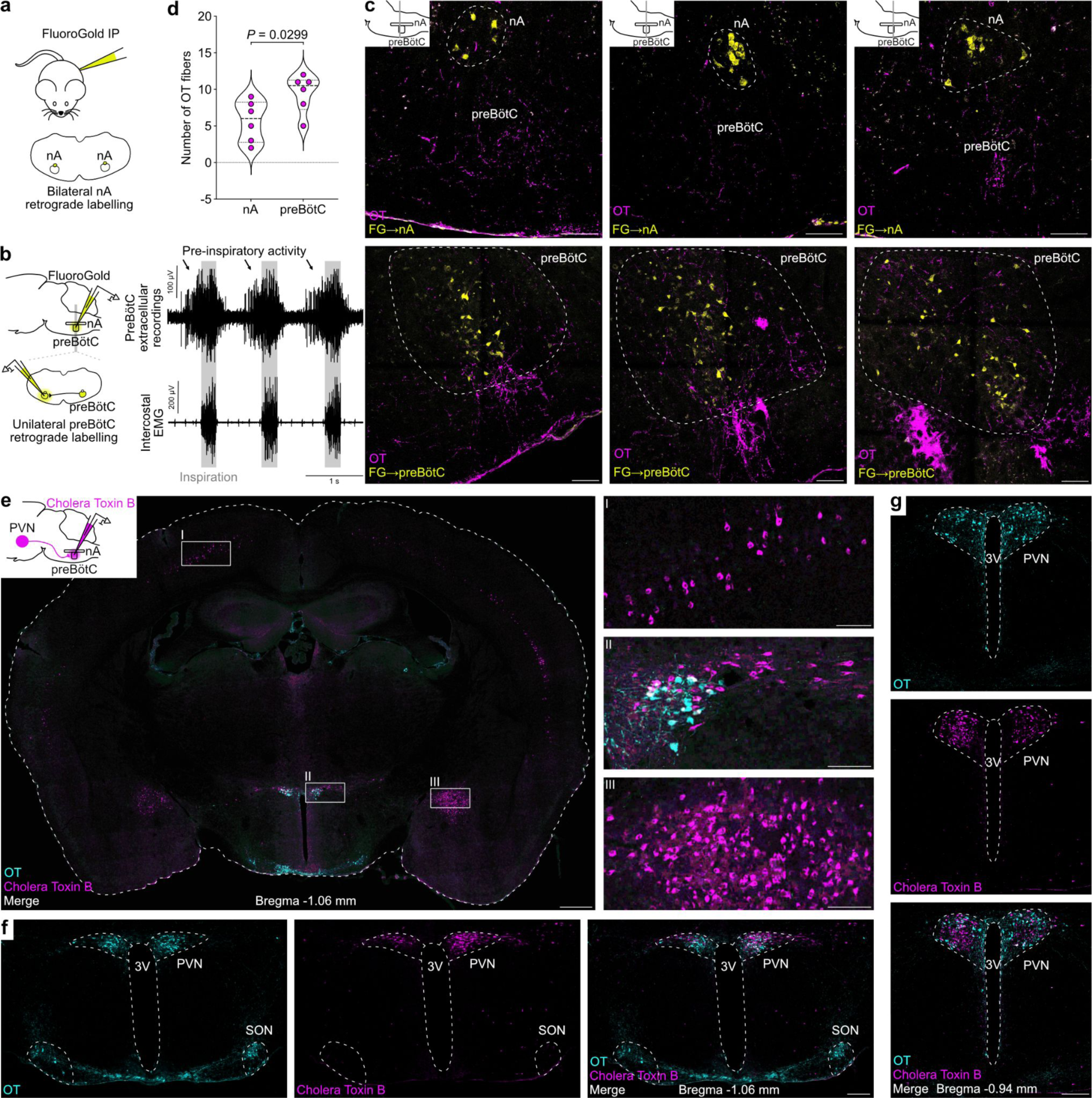
OT fibers are predominantly present in the preBötC compared to the nA, and OT neurons from the caudal PVN form a dense and regionally-specific cluster projecting to preBötC/nA neurons. **a**, Intraperitoneal (IP) injection of the retrograde tracer FluoroGold labels nucleus ambiguus neurons. **b**, FluoroGold injection in the preBötC unilaterally labels a subgroup of contralateral preBötC neurons, enabling delineation of the preBötC boundaries. PreBötC injections were guided by extracellular recordings made with the injection pipette, to localize the peak pre-inspiratory/inspiratory activity on the rostral side of the inspiratory cell column. **c**, Coronal sections at three rostro-caudal levels (insets) with immunohistochemical labeling of OT fibers, and labeling of either nA neurons (top) or contralaterally-projecting preBötC neurons (bottom) with FluoroGold (FG), using the strategies for FG injection shown in adjacent panels **a** and **b**. The preBötC contains a greater density of OT fibers compared to the nA. Scale bars, 100 µm. **d**, Each point represents the average number of OT fibers counted in three brainstem sections (50 µm thick) per animal (n = 6 for nA, n = 6 for preBötC). The sections were taken at three rostro-caudal positions through the preBötC/nA. Paired t-test for nA *vs*. preBötC. **e-g**, Bilateral injection of the retrograde tracer cholera toxin B in the preBötC/nA. OT neurons that project to the preBötC/nA are restricted to the caudal PVN. The most rostral level showing OT neurons co-labelled with cholera toxin B is at ∼ Bregma -0.94 mm (**g**). The level with the highest proportion of OT neurons co-labelled with cholera toxin B is at ∼ Bregma -1.06 mm (**e**, **f**). At this level, only three structures are labeled with cholera toxin B (**e**), the cortex (I in **e**), the PVN (II in **e**), and the central amygdala (III in **e**), similarly to what was shown previously using monosynaptic retrograde viral tracing from preBötC neurons^47^. We found no cholera toxin B labeling in the supraoptic nucleus (SON) (**f**). Scale bars, 500 µm (whole section in **e**), 100 µm (I, II and III in **e**), 200 µm (**f**, **g**).

**Extended Data Fig. 2:**
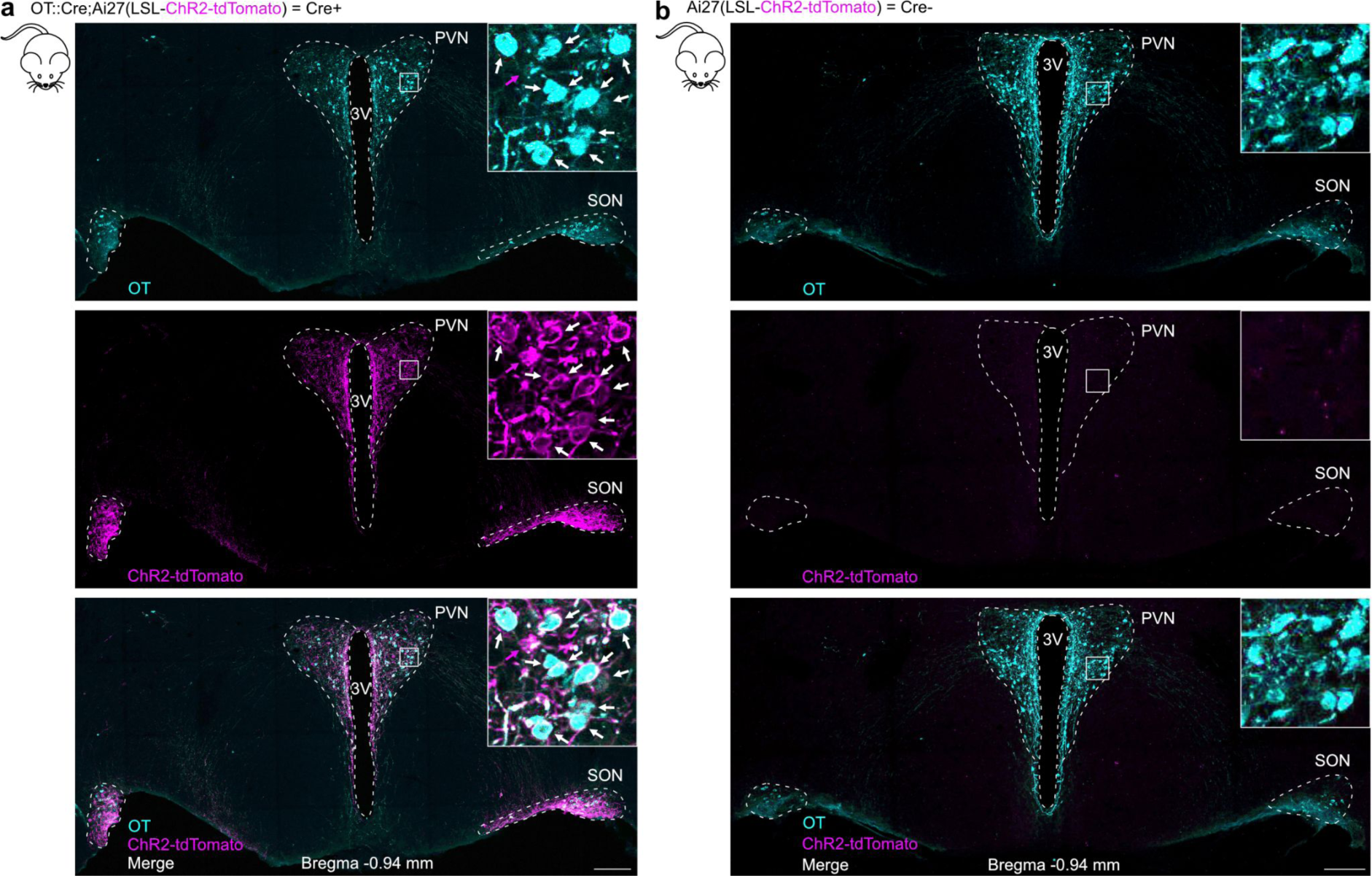
Specificity of ChR2-tdTomato expression in the OT neurons of OT::Cre;Ai27(LSL-ChR2-tdTomato) mice. **a**, The selective expression in OT neurons of the blue light gated cation channel ChR2 fused to the fluorescent protein tdTomato is induced by crossing OT::Cre mice with Ai27(LSL-ChR2-tdTomato) mice. ChR2-tdTomato labeling is mostly membranous, due to the membrane targeting of ChR2. White arrows in insets indicate neurons counted as OT^+^ and ChR2-tdTomato^+^. A magenta arrow indicates a neuron counted as OT^-^ and ChR2-tdTomato^+^, therefore representing one of the rare neurons counted as non-specific expression (3% of OT^-^ ChR2^+^ neurons, Fig. 1c). This very low non-specific expression is likely overestimated by our count, because of the limited sensitivity of OT immunohistochemical labeling leading to an underestimation of the number of OT neurons^48^, and because we counted neurons showing faint OT labeling as OT^-^ neurons (such as the one indicated by the magenta arrow). Scale bar, 200 µm. **b**, Littermates that did not express the Cre recombinase (Cre^-^), but were genotyped positive for the LSL-ChR2-tdTomato transgene, obtained from crossings between OT::Cre mice with Ai27(LSL-ChR2-tdTomato) mice, were used as control mice. No ChR2-tdTomato expression was found in these mice. Scale bar, 200 µm.

**Extended Data Fig. 3:**
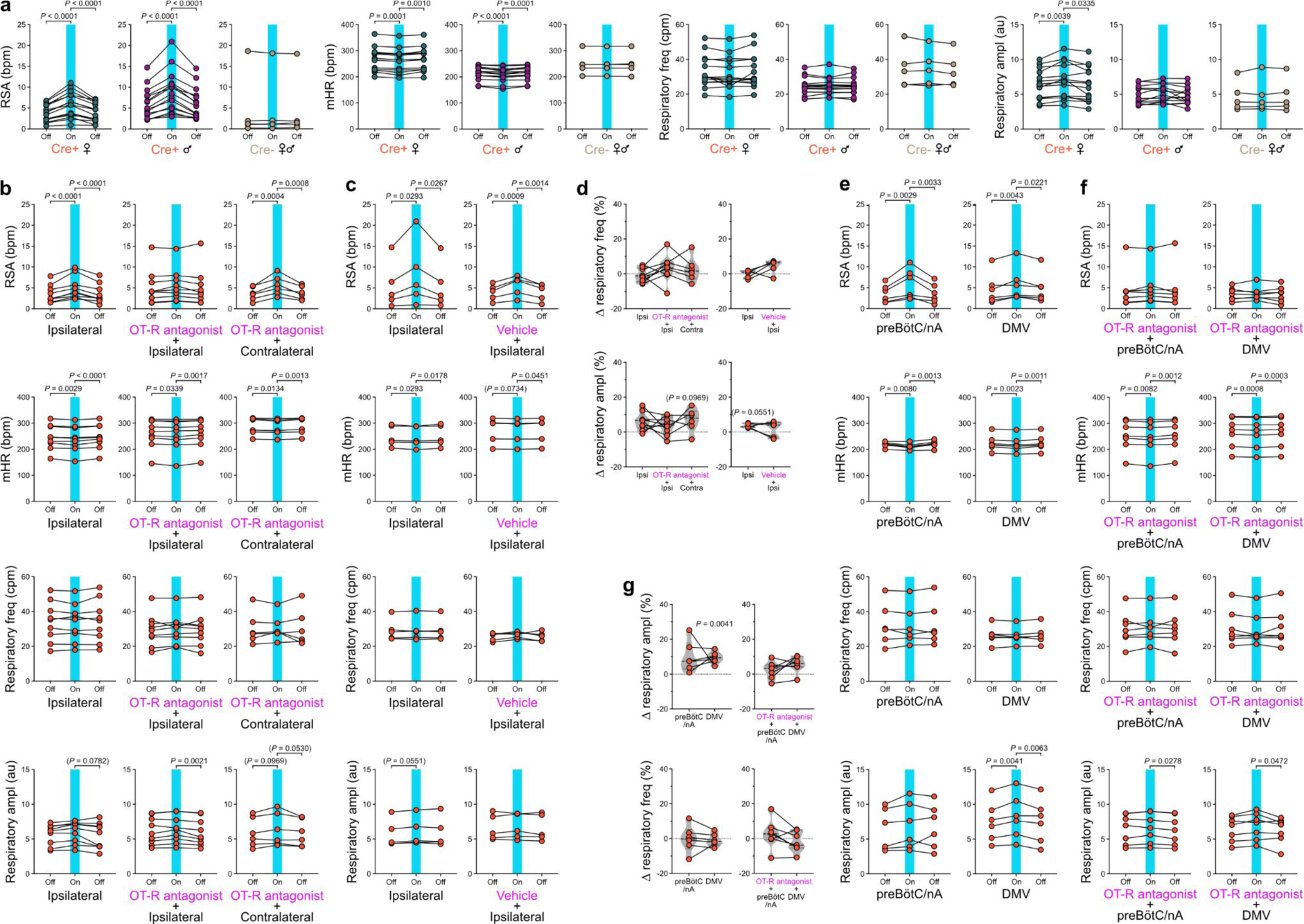
Individual data showing the respiratory and cardiac effects of the photoexcitation of PVN^OT^ fibers in the preBötC/nA and DMV, before and after injection of an OT-R antagonist in the preBötC/nA. **a**, Individual data from the photoexcitation of OT fibers unilaterally in the preBötC/nA of adult anesthetized OT::Cre;Ai27(LSL-ChR2) female and male mice (n = 15 Cre^+^ females, n = 15 Cre^+^ males, n = 5 Cre^-^ females and males). Each point is the average of ∼5-10 photoexcitation trials per mouse. RM one-way ANOVA, Tukey’s multiple comparison. The pre-photoexcitation and photoexcitation data were used to calculate the delta changes for each measured parameter represented in Fig. 2c. freq, frequency. ampl, amplitude. **b**-**c**, In a subgroup of mice shown in **a**, after ∼5-10 photoexcitation trials (“ipsilateral” data in **b** and **c**), an OT-R antagonist (**b**) or vehicle (**c**) was injected in the preBötC/nA where the photoexcitations were performed, and photoexcitations at the same ipsilateral site were repeated (“OT-R antagonist + ipsilateral” data in **b**, “vehicle + ipsilateral” in **c**) (n = 9 for “ipsilateral” and “OT-R antagonist + ipsilateral” in **b**, n = 5 for “ipsilateral” and “vehicle + ipsilateral” in **c**). In a subgroup of mice injected with the OT-R antagonist, photoexcitations in the contralateral preBötC/nA were made after the ipsilateral photoexcitations (n = 6, “OT-R antagonist + contralateral” in **b**). Each point is the average of ∼5-10 photoexcitation trials per mouse for the “ipsilateral” condition, and the average of ∼3-5 photoexcitation trials per mouse for the “OT-R antagonist + ipsilateral”, “OT-R antagonist + contralateral” and “vehicle + ipsilateral” conditions (**b**, **c**). RM one-way ANOVA, Tukey’s multiple comparison. The pre-photoexcitation and photoexcitation data were used to calculate the delta changes for the RSA and mHR parameters represented in Fig. 2g. **d**, Delta (*Δ*) changes in respiratory frequency and respiratory amplitude, from the pre-photoexcitation and photoexcitation data shown in **b** and **c**. Intra-condition delta changes (photoexcitation *vs.* pre-photoexcitation), RM one-way ANOVA, Tukey’s multiple comparison. Inter-conditions comparisons for OT-R antagonist (“ipsi” *vs.* “OT-R antagonist + ipsi” *vs.* “OT-R antagonist + contra”), RM mixed-effects analysis with the Geisser-Greenhouse correction, Tukey’s multiple comparison. Inter-conditions comparisons for vehicle (“ipsi” *vs.* “vehicle + ipsi”), paired t-test. **e**, In a subgroup of mice shown in **a**, after ∼5-10 photoexcitation trials in the preBötC/nA, photoexcitations of the ipsilateral DMV were performed (n = 6). RM one-way ANOVA, Tukey’s multiple comparison. **f**, In a subgroup of mice shown in **b**, after ∼5 photoexcitation trials in the preBötC/nA following injection of the OT-R antagonist, photoexcitations of the ipsilateral DMV were performed (n = 7). RM one-way ANOVA, Tukey’s multiple comparison. **g**, Delta changes in respiratory frequency and respiratory amplitude, from the pre-photoexcitation and photoexcitation data shown in **e** and **f**. Intra-condition delta changes (photoexcitation *vs.* pre-photoexcitation), RM one-way ANOVA, Tukey’s multiple comparison. Inter-conditions comparisons before and after OT-R antagonist injection (“preBötC/nA” *vs.* “DMV”), paired t-test.

**Extended Data Fig. 4:**
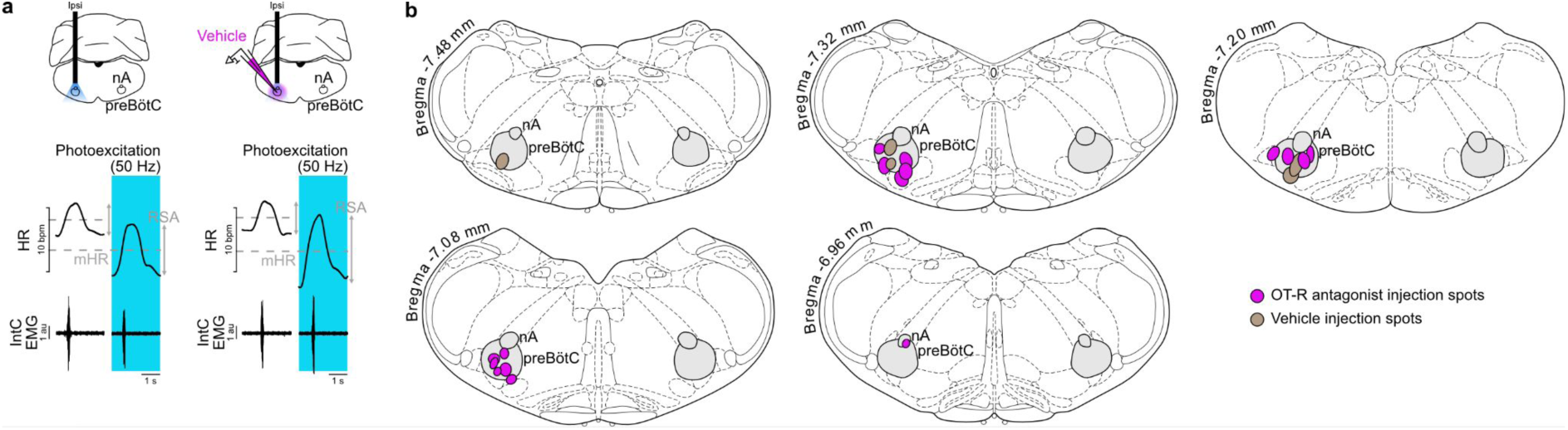
Raw traces for the photoexcitation of PVN^OT^ fibers in the preBötC/nA before and after vehicle injection, and localisation of the injection spots of the OT-R antagonist and vehicle. **a,** Vehicle injection in the preBötC/nA did not alter the effects induced by the photoexcitation of PVN^OT^ fibers (individual data in Fig. 2g and Extended Data Fig. 3c). **b**, Localisation of the injection spots of a selective OT-R antagonist (200 nl) and of vehicle (200 nl) in the preBötC/nA, for data shown in Fig. 2e-k and Extended Data Fig. 3b-g. Each spot corresponds to a single injection in one mouse.

**Extended Data Fig. 5:**
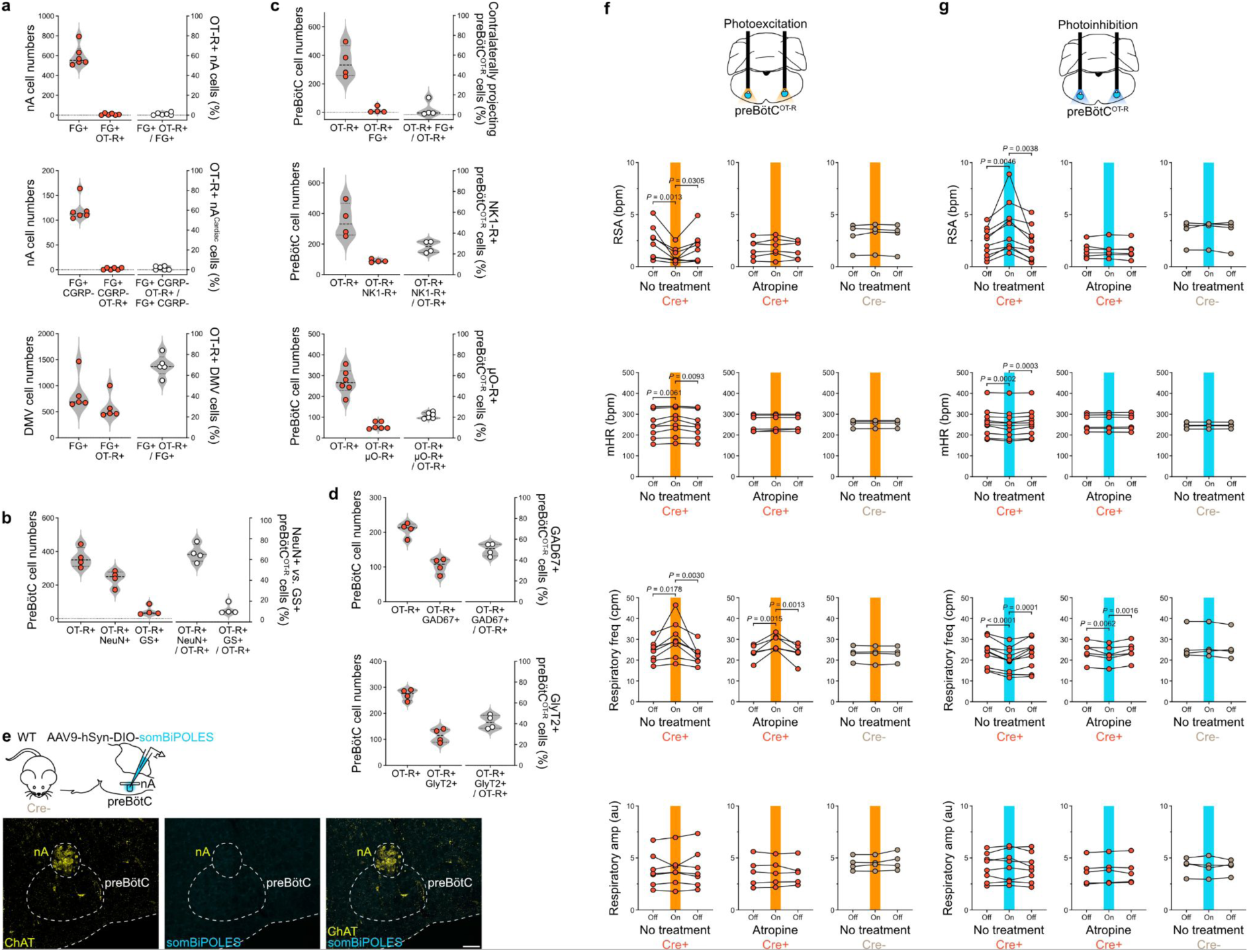
Individual data for the anatomo-functional characterisation of OT-R^+^ cells in the preBötC/nA, and absence of somBiPOLES expression in Cre^-^ mice. **a-d**, Individual data for the averaged counts shown on Fig. 3a-b and d-e. Each point represents the total number of cells counted unilaterally in one mouse (n = 6 mice for nA counts and n = 5 mice for DMV counts in (**a**); n = 4 mice in (**b**); n = 4 mice for FG^+^ counts, n = 4 mice for NK1-R^+^ counts and n = 6 mice for µO-R^+^ counts in (**c**); n = 4 mice for GAD67^+^ counts and n = 4 mice for GlyT2^+^ counts in (**d**)). **e**, Absence of somBiPOLES expression in Cre^-^ mice following viral injection in the preBötC. Scale bar, 100 µm. **f-g**, Individual data from the photoexcitation (**f**) and photoinhibition (**g**) of preBötC^OT-R^ neurons bilaterally in adult anesthetized mice (n = 8 Cre^+^ mice for photoexcitation and n = 10 Cre^+^ mice for photoinhibition with no treatment, n = 6 Cre^+^ mice for photoexcitation/photoinhibition after atropine, n = 4 Cre^-^ mice). Each point corresponds to the average of ∼5 photoexcitation or photoinhibition trials per mouse. RM one-way ANOVA, Tukey’s multiple comparison. The pre-photomodulation and photomodulation data were used to calculate the delta changes for each measured parameter represented in Fig. 3h and i. freq, frequency. ampl, amplitude.

**Extended Data Fig. 6:**
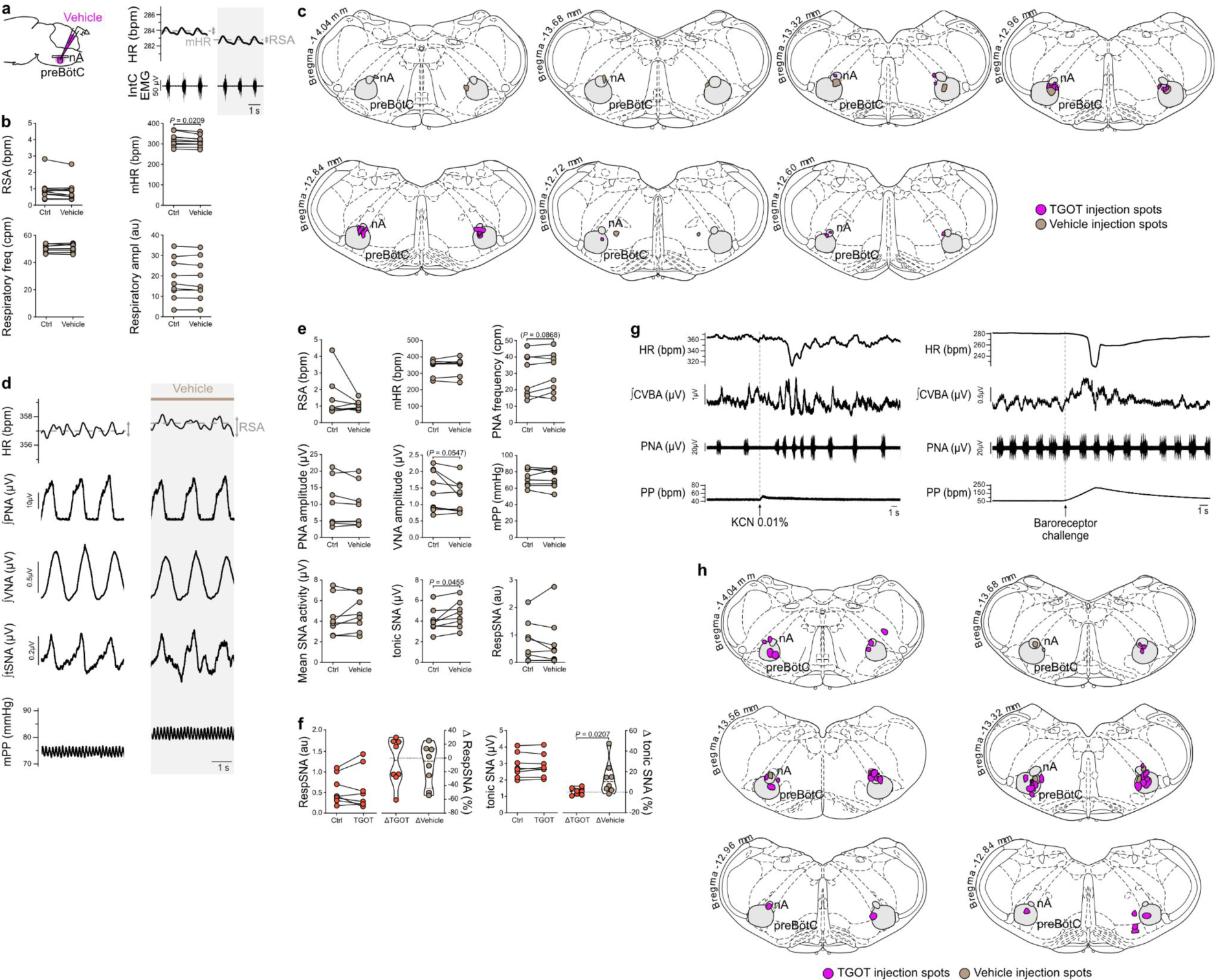
Vehicle injections in the preBötC of adult anesthetized rats and *in situ* Working Heart-Brainstem Preparations of juvenile rats, maps of preBötC injections, and functional characterisation of CVBA recordings. **a-b**, Bilateral injection of vehicle in the preBötC of adult anesthetized Wistar rats had no effect on HR and inspiratory activity (IntC EMG). Absolute values are represented for the conditions before and after vehicle injection (n = 10). Paired t-test for respiratory frequency, respiratory amplitude and mHR, Wilcoxon matched-pairs signed rank test for RSA. **c**, Schematic coronal maps showing the distribution of all TGOT and vehicle injection spots in adult anesthetized Wistar rats. **d-e**, Effects of vehicle injection in the preBötC of *in situ* Working Heart-Brainstem Preparations of juvenile Wistar rats (P21-30) (HR, n = 8; PNA frequency, n = 9; PNA amplitude, n = 8; VNA, n = 9; tSNA, n = 8; mPP, n = 8). Integrated traces (**d**) are respiratory triggered averages over >15 respiratory cycles. Absolute values are represented for the conditions before and after vehicle injection (**e**). Paired t-test or Wilcoxon matched-pairs signed rank test. **f**, Effects of bilateral TGOT *vs.* vehicle injection in the preBötC of *in situ* Working Heart-Brainstem Preparations of juvenile Wistar rats on the two components of tSNA, the phasic burst of respiratory modulation of tSNA (RespSNA) and the tonic SNA (n = 8 TGOT and n = 8 vehicle). Before vs. after injection conditions, paired t-test. Comparison of the relative effects of TGOT and vehicle (delta (*Δ*) in %), unpaired t-test for RespSNA, Mann-Whitney test for tonic SNA. **g**, Peripheral chemoreflex (potassium cyanide (KCN) injection) and baroreflex (increased perfusion flow rate) challenges were used to confirm functionally the validity of the CVBA recordings (data shown from the same preparation). **h**, Schematic coronal maps showing the distribution of all TGOT and vehicle injection spots in the *in situ* Working Heart-Brainstem Preparations of juvenile Wistar rats.

**Extended Data Fig. 7:**
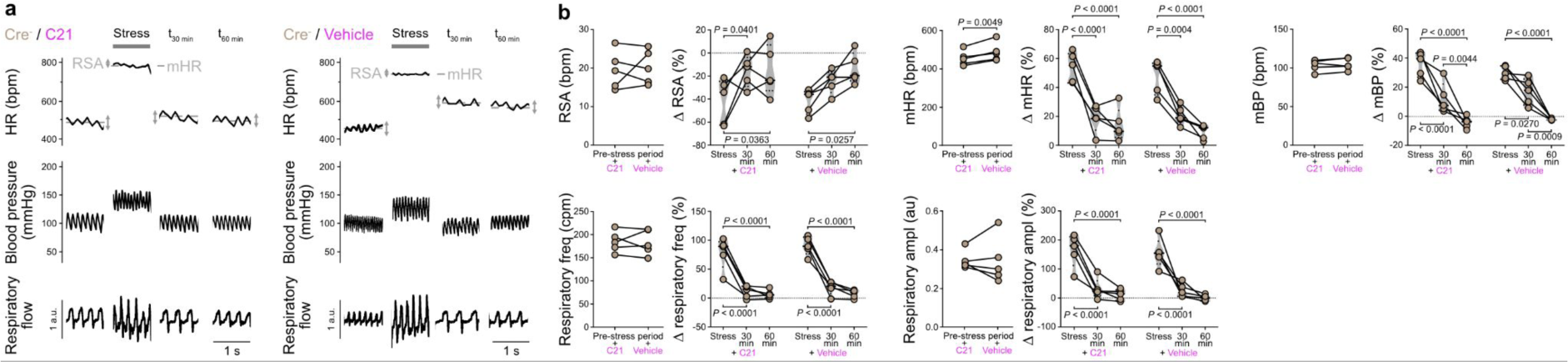
The DREADD agonist Compound 21 is mostly inert on cardiovascular and respiratory functions in OT::Cre;R26-LSL-hM4Di-DREADD Cre^-^ mice, except for mHR. **a-b**, Effects induced by the intraperitoneal injection of the hM4Di agonist Compound 21 (C21) or vehicle, before, during and after a restraint stress test in a ventilated cylinder, in Cre^-^ mice from the OT::Cre;R26-LSL-hM4Di-DREADD transgenic mouse line (n = 7, three trials averaged for each condition, C21 and vehicle, in each mouse; C21 or vehicle trials spaced by at least 24h). Traces on the right for both the C21 and vehicle conditions are enlargements of the gray shaded areas on their respective traces on the left (**a**). There were no off-target effects of C21 in Cre^-^ mice on RSA amplitude, mBP, respiratory frequency and respiratory amplitude, but mHR which was increased during the pre-stress condition. Pre-stress conditions, C21 *vs.* vehicle, paired t-test (**b**). Stress and post-stress delta (*Δ* in %) effects compared to the pre-stress condition, C21 *vs.* vehicle, RM two-way ANOVA, Sidak’s multiple comparison.

**Extended Data Fig. 8:**
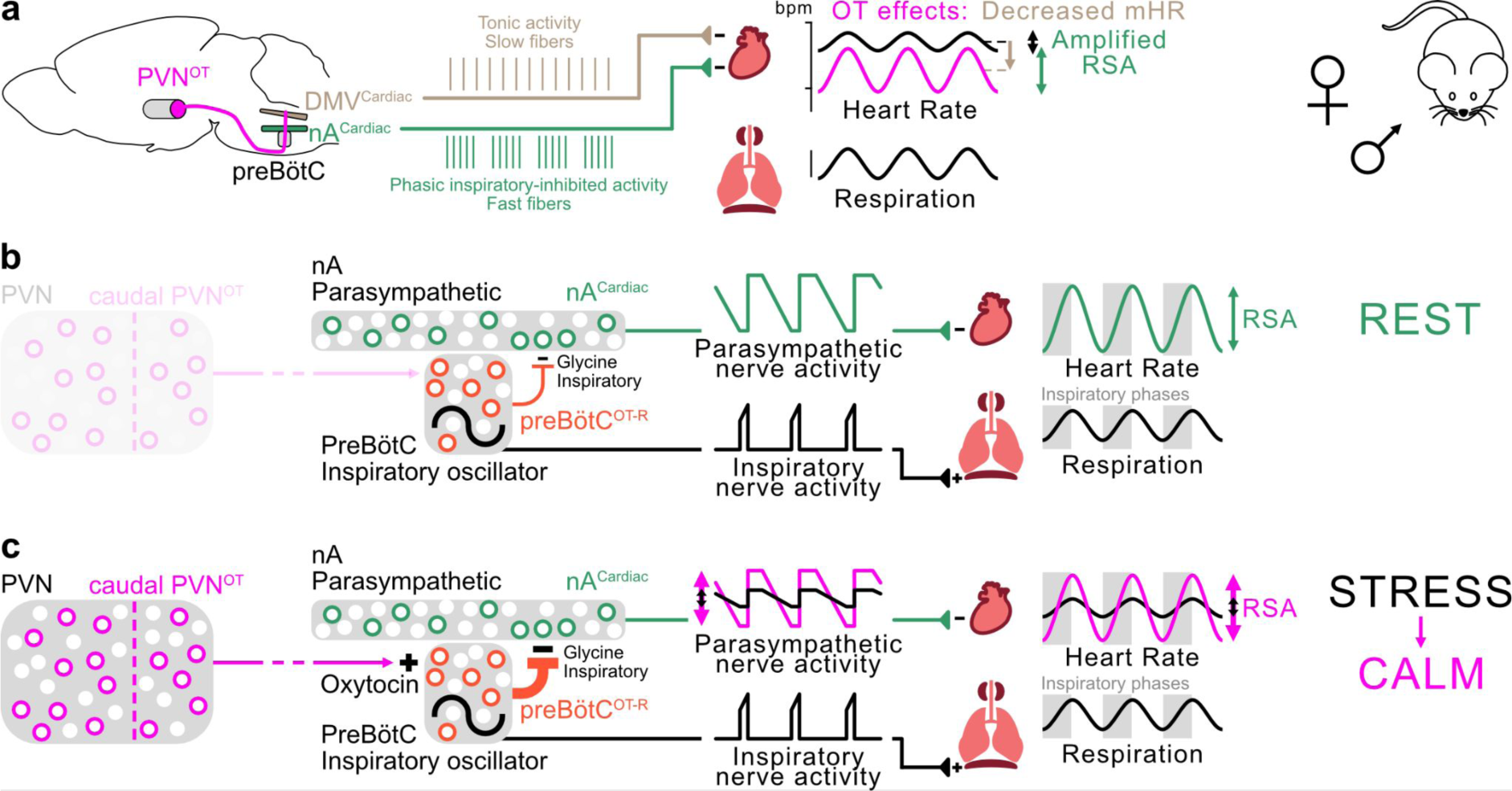
Schematic model of the preBötC^OT-R^→nA^Cardiac^ neuronal circuit that controls RSA at rest, and its modulation by PVN^OT^ neurons for RSA amplification during a calming behavior. **a**, Oxytocinergic neurons from the caudal PVN project to the brainstem, with axons traversing the medulla oblongata in the coronal plane and crossing cardiorespiratory nuclei including the preBötC, nA and DMV. The stimulation of these fibers releases OT in the preBötC/nA which induces RSA amplification, and concomitantly induces a mHR decrease *via* the DMV. The OTergic amplification of RSA was found in both female and male mice and rats. **b**, At rest, OT is not involved in the regulation of RSA amplitude. Yet, expression of the OT-R in a subgroup of preBötC neurons, the group that generates the inspiratory rhythm, defines a subpopulation of cells that participates in the control of RSA. PreBötC^OT-R^ neurons induce inspiratory glycinergic inhibition of the activity of nA^Cardiac^ neurons *via* putative monosynaptic contacts. This results in a decrease in cardiac parasympathetic activity that is concomitant with each inspiratory burst in the phrenic nerve, and at the effector level in an increase in HR during inspiration. **c**, PVN^OT^ neurons are involved in the recovery of RSA amplitude during calming behavior following a stress. The release of OT by PVN^OT^ neurons on preBötC^OT-R^ neurons amplifies RSA, by raising the excitability of inspiratory preBötC neurons leading to the amplification of the preBötC→nA^Cardiac^ inspiratory glycinergic connectivity. The RSA amplification is generated *via* amplified respiratory modulation of parasympathetic activity to the heart. Inhibition of PVN^OT^ neurons slows the recovery of RSA amplitude following a stress.

